# Disproportionate CH_4_ sink strength from an endemic, sub-alpine Australian soil microbial community

**DOI:** 10.1101/2020.11.08.373464

**Authors:** M.D. McDaniel, M. Hernández, M.G. Dumont, L.J. Ingram, M.A. Adams

## Abstract

Soil-to-atmosphere methane (CH_4_) fluxes are dependent on opposing microbial processes of production and consumption. Here we use a soil-vegetation gradient in an Australian sub-alpine ecosystem to examine links between composition of soil microbial communities, and the fluxes of greenhouse gases they regulate. For each soil-vegetation type (forest, grassland, and bog), we measured carbon dioxide (CO_2_) and CH_4_ fluxes and their production/consumption at 5-cm intervals to a depth of 30 cm. All soils were sources of CO_2_, ranging from 49-93 mg CO_2_ m^-2^ h^-1^. Forest soils were strong net sinks for CH_4_ at rates up to −413 µg CH_4_ m^-2^ h^-1^. Grassland soils varied with some soils acting as sources and some as sinks, but overall averaged −97 µg CH_4_ m^-2^ h^-1^. Bog soils were net sources of CH_4_ (+340 µg CH_4_ m^-2^ h^-1^). Methanotrophs were dominated by USCα in forest and grassland soils, and *Candidatus* Methylomirabilis sp. in the bog soils. *Methylocystis* were also detected at relatively low abundance. The potential disproportionately large contribution of these ecosystems to global CH_4_ oxidation, and poorly understood microbial community regulating it, highlight our dependence on soil ecosystem services in remote locations can be driven by a unique population of soil microbes.

**Originality-Significance Statement:** (Identify the key aspects of originality and significance that place the work within the top 10% of current research in environmental microbiology)

Novel methanotrophic bacteria have been discovered in recent years, but few studies have examined the total known diversity of methanotrophs together with the net flux of CH_4_ from soils. We used an ecosystem with a vegetation-soil gradient in the sub-alpine regions of Australia (with extremely strong consumption of atmospheric CH_4_) to examine microbial and abiotic drivers of CH_4_ fluxes across this gradient. Recently characterized methanotrophs, either USCα in forest and grassland soils, or oxygenic *Candidatus* Methylomirabilis sp. in the bog soil were dominant. Methanotrophs belonging to the families Methylococcaceae and Methylocystaceae represented only a small minority of the methanotrophs in this ecosystem.

## Introduction

Counteracting biogeochemical processes that consume or produce greenhouse gases (GHGs) regulate whether soils act as net sources or sinks. The magnitude and spatial distributions of these competing processes – whether across landscapes, with soil depth (Conrad and Rothfuss, 1991; Bender and Conrad, 1994; He *et al*., 2012), or even within soil aggregates (Sexstone *et al*., 1985; Ebrahimi and Or, 2016; Karbin *et al*., 2016) – determine net release or uptake. Soils, especially upland and older soils, are nearly always sources of carbon dioxide (CO_2_) to the atmosphere, owing to production of CO_2_ via heterotrophic and root respiration, which overwhelm slow rates of autotrophic CO_2_ consumption. On the other hand, soils can routinely be either sources or sinks for methane (CH_4_) and nitrous oxide. In some cases, soils can switch from being a net source to a net sink for CH_4_ and N_2_O in a matter of minutes to hours (Harriss *et al*., 1982; Wille *et al*., 2008; McDaniel *et al*., 2014; Hernández *et al*., 2017). Similarly, a lateral distance of just a meter or two may result in sinks becoming sources, or vice versa (Velthof *et al*., 1996; Priemé *et al*., 1997; McDaniel *et al*., 2017).

Methane is a potent greenhouse gas, 34 times as potent as CO_2_, and is responsible for ∼17% of anthropogenic warming (Stocker, 2013). Soil sources and sinks for CH_4_ are controlled by abundance and composition of specific microbial communities (Murrell and Jetten, 2009; Nazaries, Murrell, *et al*., 2013), but also regulated by abiotic factors (Sullivan *et al*., 2010; Wu *et al*., 2010; Wolf *et al*., 2012). Oxidation of CH_4_ to microbial biomass and CO_2_ is, for example, restricted to distinct groups of specialized methanotrophic microorganisms. Aerobic methanotrophs belonging to the α-Proteobacteria and δ-Proteobacteria have been studied for decades and detected across a wide range of habitats (Knief, 2015). Groundbreaking research in recent years has uncovered previously uncharacterized methanotrophs, including acidophilic Verrucomicrobia from the family Methylacidiphilaceae (Khadem *et al*., 2012) and intra-oxygenic methanotrophs, *candidatus* Methylomirabilis oxyfera, from the NC10 phylum (Ettwig *et al*., 2010). In addition, more recent studies have identified USCα methanotrophs, which are specialized atmospheric-CH_4_ consuming bacteria whose activity has been known for decades but had evaded efforts to be isolated or characterized (Pratscher *et al*., 2018; Singleton *et al*., 2018; Tveit *et al*., 2019). The first step in the biochemical pathways of CH_4_ oxidation is catalyzed by the methane monooxygenase (MMO) enzyme. The *pmoA* gene encoding a subunit of the membrane-bound MMO (pMMO) enzyme is present in most aerobic methanotrophs and is frequently used as a genetic marker.

Methanogenesis in soil typically requires anaerobic and low redox conditions, with depletion of strong electronegative terminal electron acceptors, e.g. nitrate, iron. Methanogens belong to the Euryarchaeota phylum, and produce CH_4_ either from CO_2_ and H_2_ (hydrogenotrophic methanogenesis), acetate (acetoclastic methanogenesis), or methylated single-C one-carbon compounds (Thauer et al., 2008). All known methanogenic pathways include the methyl-coenzyme M reductase enzyme encoding the *mcrA* gene, which is used as a genetic marker for methanogens in the environment. Most aerobic soils show little production of CH_4_, as evidenced by low abundance of *mcrA* genes (Hernández *et al*., 2017). A few recent studies have shown that even water-logged soil profiles tend to have an aerobic upper layer that, via its capacity for CH_4_ oxidation, helps mitigate CH_4_ release to the atmosphere from lower soil depths. In some cases this layer can remove 80-97% of CH_4_ produced at lower depths (Conrad and Rothfuss, 1991; Frenzel *et al*., 1992; Oremland and Culbertson, 1992; Sass *et al*., 1992). In the field, and at ecosystem spatial scales, our understanding of the drivers of soil GHG fluxes is profoundly limited.

Local soil and vegetation gradients provide an opportunity to explore mechanisms that regulate soil GHG fluxes (Freitag *et al*., 2010; Krause *et al*., 2013; Christiansen *et al*., 2016). Soil biogeochemical conditions (e.g. microbial substrates, reduction-oxidation potential, and soil pH) can vary dramatically over relatively short distances despite more-or-less constant climate. We used a forest-grassland-bog gradient located within a sub-alpine region of southern New South Wales (Fig. 1), Australia to examine the drivers of variation in soil microbial communities, and CO_2_ and CH_4_ fluxes. These soils are of particular interest given their previously-observed, rapid rates of CH_4_ oxidation in forest and grassland soils and likely biogeochemical sensitivity to climate change. Our objective was to determine the role of soil microbial communities in regulating the production/consumption of CH_4_ across this soil-vegetation gradient.

**Fig. 1.**
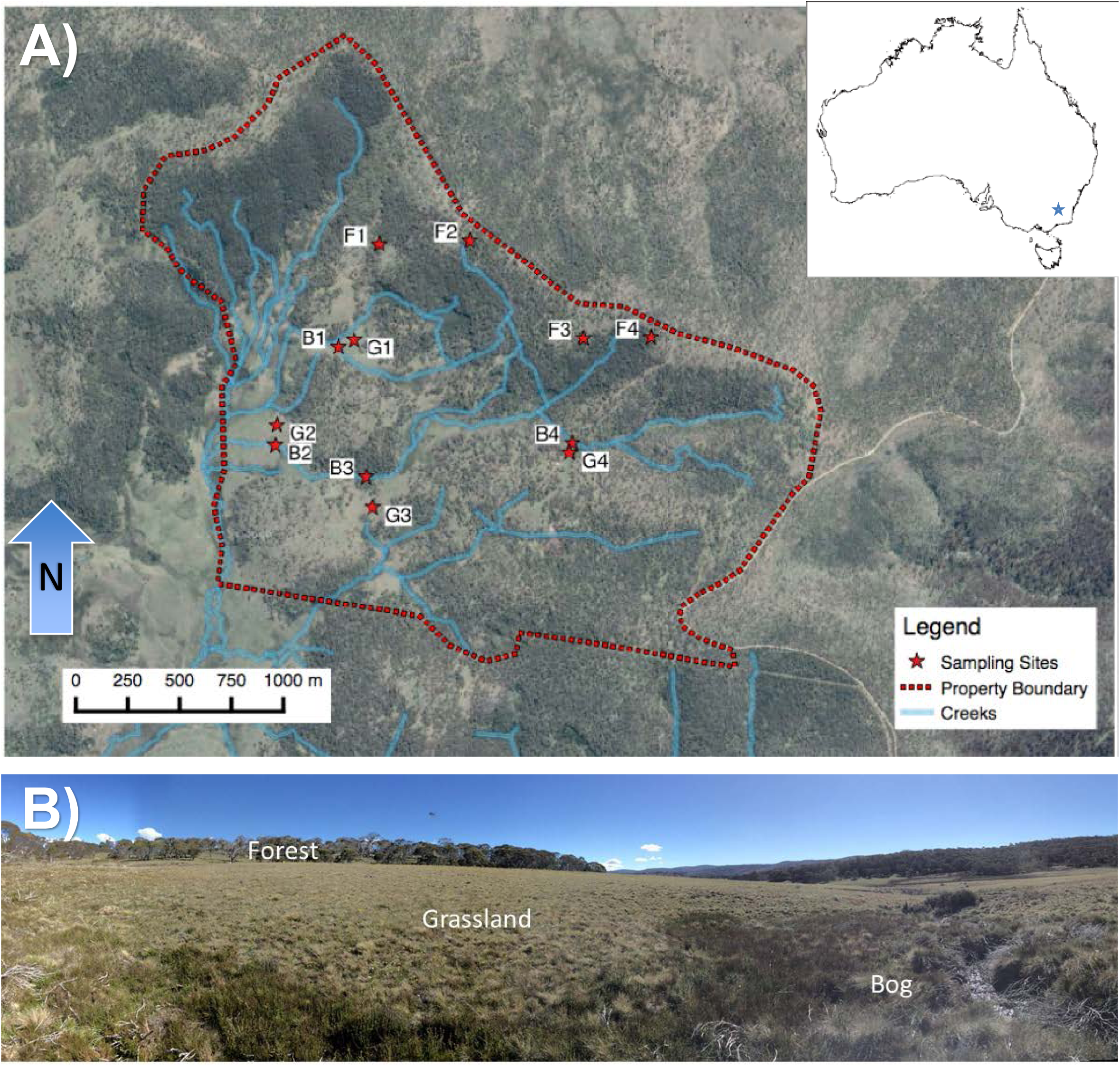
**A)** Location of 548 ha experiment area (inset of Australia), sampling sites within area and nearby streams. **B)** Landscape-level photograph of the vegetation gradient from sphagnum-dominated bog in foreground to eucalyptus-dominated forest in background. Abbreviations are F = Forest, G = Grassland, and B = Bog. The numbers after the letter represent which transect the sampling location belongs to.

## Results

### Soil greenhouse gas fluxes and production at depth

Surface fluxes of CO_2_ and CH_4_ were measured once each season. Air temperatures ranged from 3.1 to 28.3 °C, while the average soil temperature at 0-7 cm depth was 3.6 to 16.4 (Table S1). Gravimetric water content on 17 February, when microbial community analyses were conducted, ranged from 0.25-0.79 for forest soils, 0.25-0.38 for grassland soils, and 0.9-4.66 g H_2_O g dry soil^-1^ for bog soils (Table S2). The relative differences in soil temperature and moisture amongst soil-vegetation types during February extend to other three sampling dates (Table S1,S2).

**Table 1.**
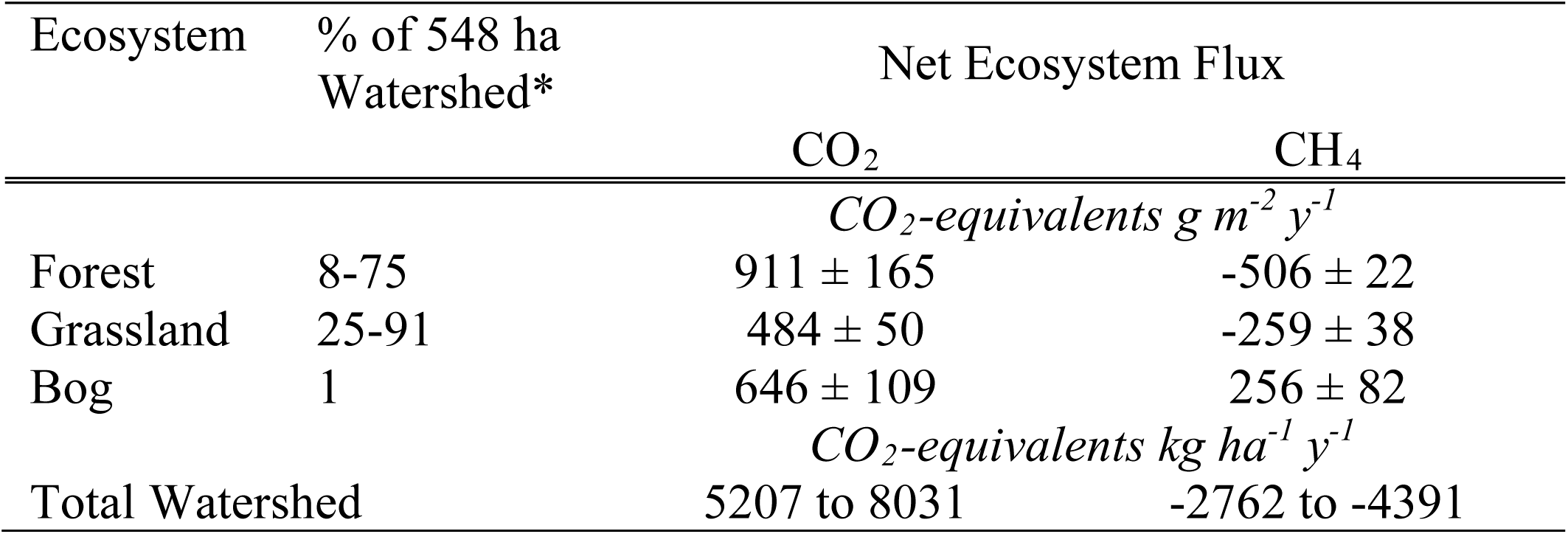
Annual estimates (mean, standard error, and range) for net ecosystem flux of carbon dioxide and methane

**Table 2.** Comparison of Global and Australian Alps soil methane sink estimates

| Measurement | Values |  |  |
| --- | --- | --- | --- |
| *Ecosystems Areal Coverage* |  |  |  |
| Global forest+grassland ecosystems (millions of ha) | 5,100 |  |  |
| Areal coverage of Australian Alps – Figure 8 (millions of ha) | 1.23 |  |  |
| Fraction of Australian Alps to global forest+grassland (%) | 0.024 |  |  |
| *Soil CH<sub>4</sub> Sink Estimates* |  |  |  |
|  | **Low**<br>**Estimate** | **Best**<br>**Estimate** | **High**<br>**Estimate** |
| Global CH<sub>4</sub> Soil Sink<sup>†</sup> (Tg y<sup>-1</sup>) | -9 | -30 | -100 |
| Mean annual Australian Alps CH<sub>4</sub> sink<sup>‡</sup> (kg ha<sup>-1</sup> y<sup>-1</sup>) | -4.2 | -19.2 | -33.2 |
| *Australian Alps Contribution to Global CH<sub>4</sub> Sink (%)* |  |  |  |
|  | **Low**<br>**Estimate** | **Best**<br>**Estimate** | **High**<br>**Estimate** |
| Low estimate (-6.2 kg ha<sup>-1</sup> y<sup>-1</sup>) | 69 | 21 | 6 |
| Best Estimate, or our projection (-19.2 kg ha<sup>-1</sup> y<sup>-1</sup>) | 213 | 64 | 19 |
| High estimate (-32.3 kg ha<sup>-1</sup> y<sup>-1</sup>) | 359 | 108 | 32 |

†: Low and Best Estimates from Kirschke et al. (2013) and Saunois et al. (2016). Other estimates have a high estimates of -100 Tg y<sup>-1</sup> (Smith et al. 2000)
‡: Based on forest foliage imagery, soil temperatures and moisture estimates, and CH<sub>4</sub> modeling described in Experimental Procedures. High and Low estimate from +/- relative standard deviation from ecosystem means.

Mean soil-to-atmosphere CO_2_ fluxes (measured *in situ*) across the soils ranged from 2.5 to 17.4 mg CO_2_ m^-2^ h^-1^, spring and summer respectively (Fig. 2A-D). Belowground, CO_2_ production (measured in the laboratory) decreased with depth. CO_2_ production was predominately from top 0-5 cm of soil, with soil means ranging from 72 to 2,357 µg CO_2_ g^-1^ d^-1^. Whereas at the 25-30 cm depth means ranged from 4 to 197 µg CO_2_ g^-1^ d^-1^. Forest and bog soils showed significantly greater CO_2_ production in summer than the grassland soil at multiple depths (*P* < 0.01, Fig. 2E), ranging from 47 to 398% greater in forest soil or 60 to 282% greater in bog soil. Whereas, there were less pronounced differences amongst soil types in autumn, winter, and spring.

**Fig. 2.**
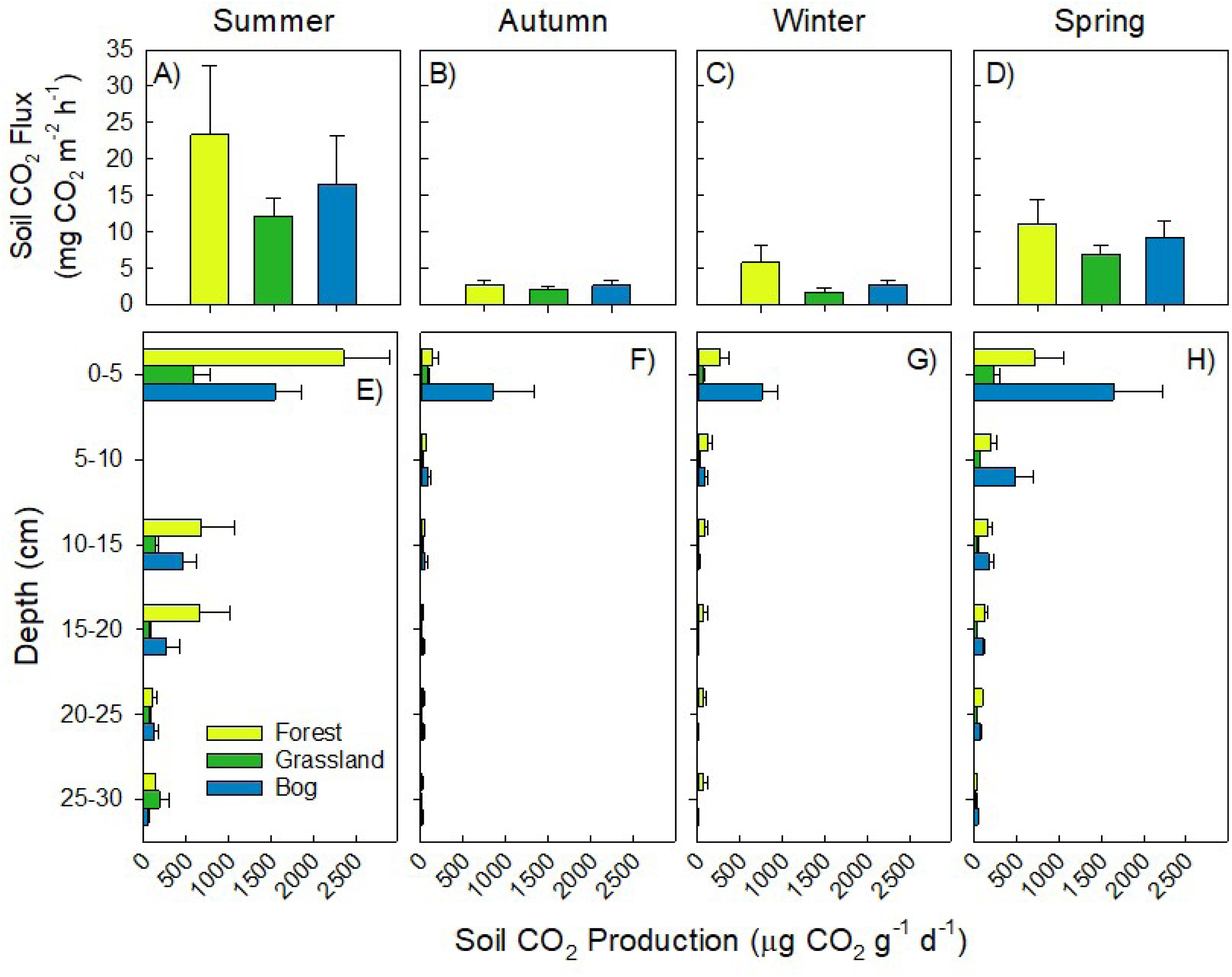
Soil CO_2_ fluxes (top panels, **A-D**) and production (bottom panels, **E-H**). Surface flux measurements and soils collected for production on 17 February (Summer,**A** & **E**), 25 May (Autumn, **B** & **F**), 22 September (Winter, **C** & **G**), and 23 November (Spring, **D** & **H**) in 2015. Mean and standard error shown (n = 4).

*In situ* CH_4_ fluxes ranged from −413 to +778 µg CH_4_ m^-2^ h^-1^ (Fig. 3A-D). Absolute fluxes, whether positive or negative, were greatest in summer compared to other seasons. Although similar pattern of soil type persisted throughout other three seasons. Forest and grassland soils mostly consumed CH_4_ throughout the year (Fig. 3A-D), while bog soils were net producers of CH_4_ in three of four seasons (Fig. 3A-C). Belowground, most soils (including bog) showed net CH_4_ consumption at all depths to 30 cm (Fig. 3E-H). The main exception was summer where large CH_4_ production of 3838 µg CO_2_ g^-1^ d^-1^ was found at 5-10 cm depth (*P* = 0.003), as well as moderate production of CH_4_ by all soils at 25-30 depth ranging from 87-704 µg CO_2_ g^-1^ d^-1^ (Fig. 3E). The net global warming potential, measured as CO_2_-equivalents, provides further evidence as the exceptional strength of the soil CH_4_ sink in this ecosystem (Table 1).

**Fig. 3.**
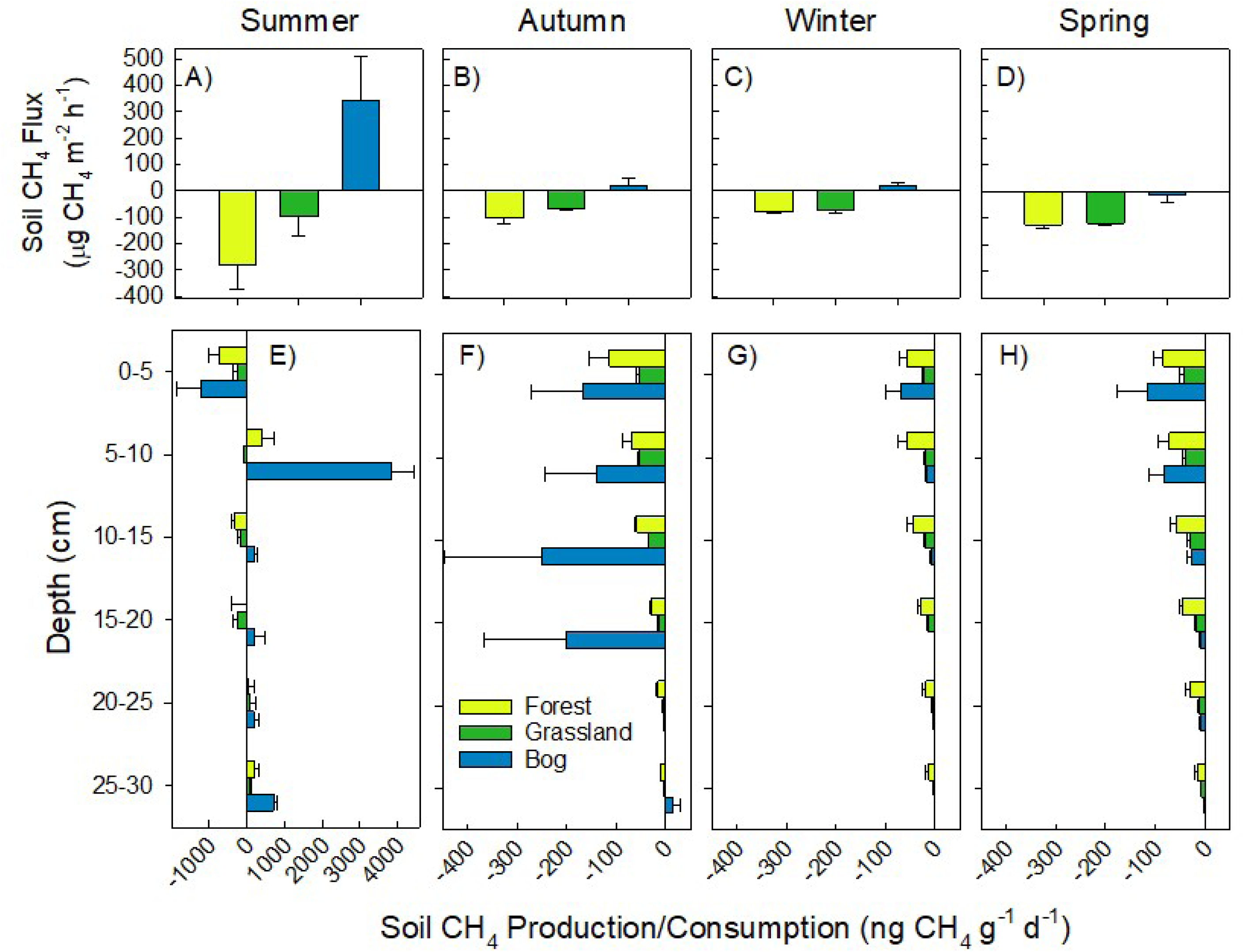
Soil CH_4_ fluxes (top panels, **A-D**) and production/consumption (bottom panels, **E-H**). Surface flux measurements and soils collected for production on 17 February (Summer,A & E), 25 May (Autumn, **B** & F), 22 September (Winter, **C** & **G**), and 23 November (Spring, **D** & **H**) in 2015. Mean and standard error shown (n = 4).

### Archaeal and bacterial gene abundance (qPCR) during summer (17 February 2015)

There were significant differences in archaeal 16S rRNA gene abundance between soil types, with the bog soil showing one to two orders of magnitude greater abundances than the other two soils at multiple depths (*P* < 0.060, Fig. 4A). The abundances of bacterial 16S rRNA genes were highly variable but did show a decrease with depth from 2.3 × 10^11^ to 1.6 × 10^10^ copies per g soil (Fig. 4B). There were no significant differences in abundance of bacterial 16S rRNA genes among soil types. A *pmoA* qPCR assay targeting conventional methanotrophs belonging to the families Methylococcaceae and Methylocystaceae showed highest abundance in the bog samples (Fig 4C).

**Fig. 4.**
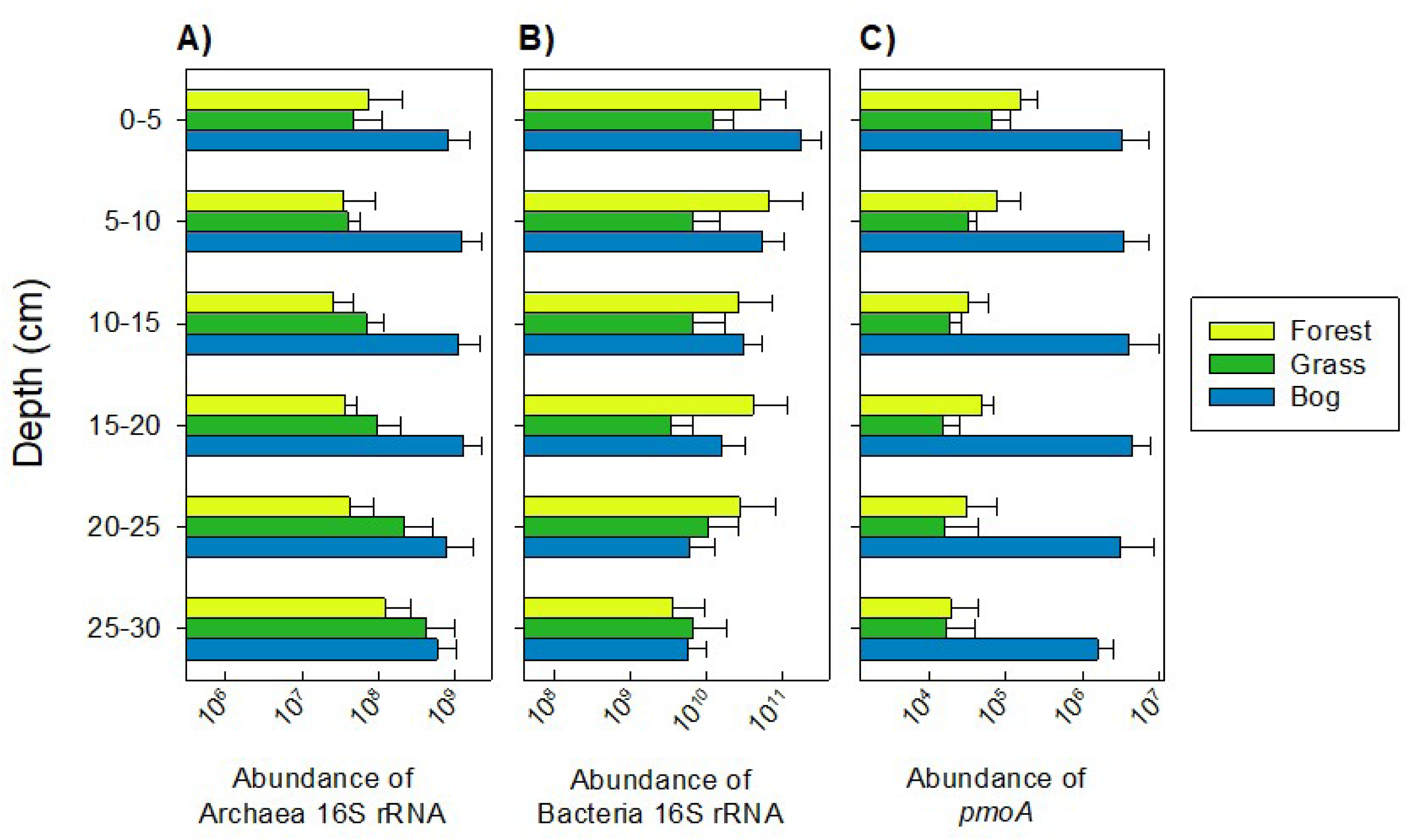
Abundance of archaeal (**A**) and bacterial (**B**) 16S rRNA genes, and *pmoA* (C). Means and standard error are shown (n=4) for all samples.

Across the soil-vegetation gradient, GHG production at depth was positively related to the soil-to-atmosphere flux of both gases (Fig. 5A, B). There was also evidence for GHG production correlating with specific gene abundances across all soil types and depths. For example, the abundance of bacterial 16S rRNA genes was linearly, positively related to CO_2_ production (Fig. 5C). While, the abundance of archaeal 16S rRNA was non-linearly, positively related to CH_4_ production (3-parameter exponential equation, Fig. 5D). Interestingly, however, we did not find significant relationships between abundances of archaea and CO_2_, nor between bacterial abundances and CH_4_.

**Fig. 5.**
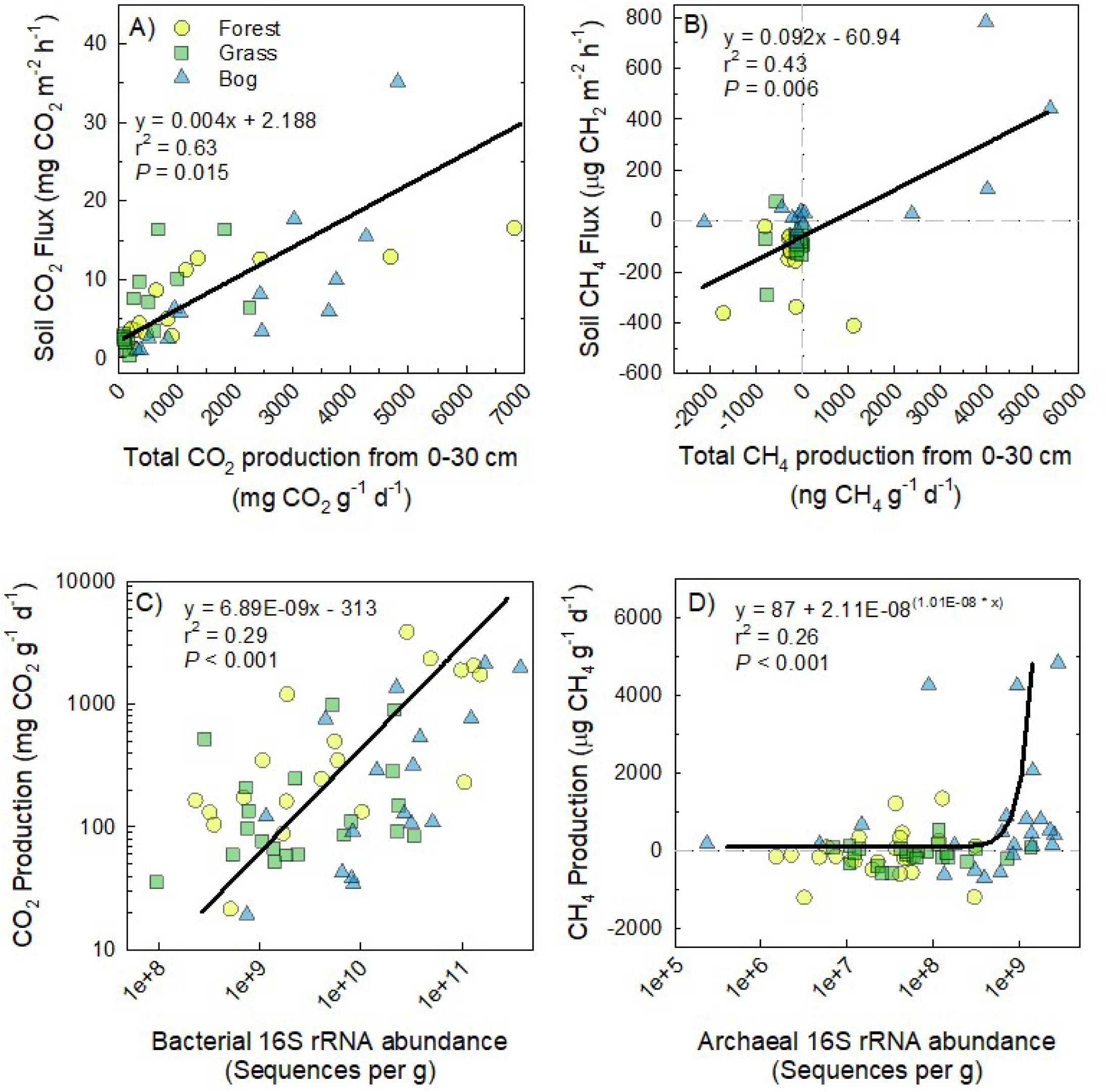
Regressions between total CO_2_ production and CO_2_ soil-atmosphere flux (**A**), CH_4_ production and CH_4_ soil-atmosphere flux (**B**). Total GHG production was calculated for each soil profile as sum of production from 0 to 30 cm depth. Regressions between bacterial 16S rRNA abundance with CO_2_ production (**C**, linear equation with log, log scale). Regression between and archaeal 16S rRNA and CH_4_ production (**D**, 3-parameter exponential, log x-axis). Equations, chosen by best fit, are shown in each panel.

### Microbial community composition and diversity during summer (17 February 2015)

Across all soils, Illumina sequencing resulted in 440,504 archaeal, 24,293,004 bacterial, and 2,258,405 *pmoA* sequences. Both clustering and multivariate analyses (Fig. 6) show distinct differences in bacterial 16S rRNA genes between the forest/grassland and bog soils, with less forest and grassland soils being more similar. In the bog soils, *Betaproteobacteria* (OTU 1850) and OTU-1522 were more abundant than in forest and grassland soils (Fig. 6A). Contrastingly, *Rhizobiales* (OTU-78) and OTU-71 differentiated the forest/grassland from bog soils. A few OTUs were uniquely abundant in the grassland soils, like the *Chloroflexi* group (especially OTU-252, OTU-1043, OTU-249). Across multiple depths, the bog soils had greatest 16S rRNA diversity compared to the forest and grassland soils (Fig. S1). The bog soils had greater diversity, 10% and 42% larger Shannon diversity and richness (H’ and S), than the average of forest and grassland soils across all depths (*P* < 0.001).

**Fig. 6.**
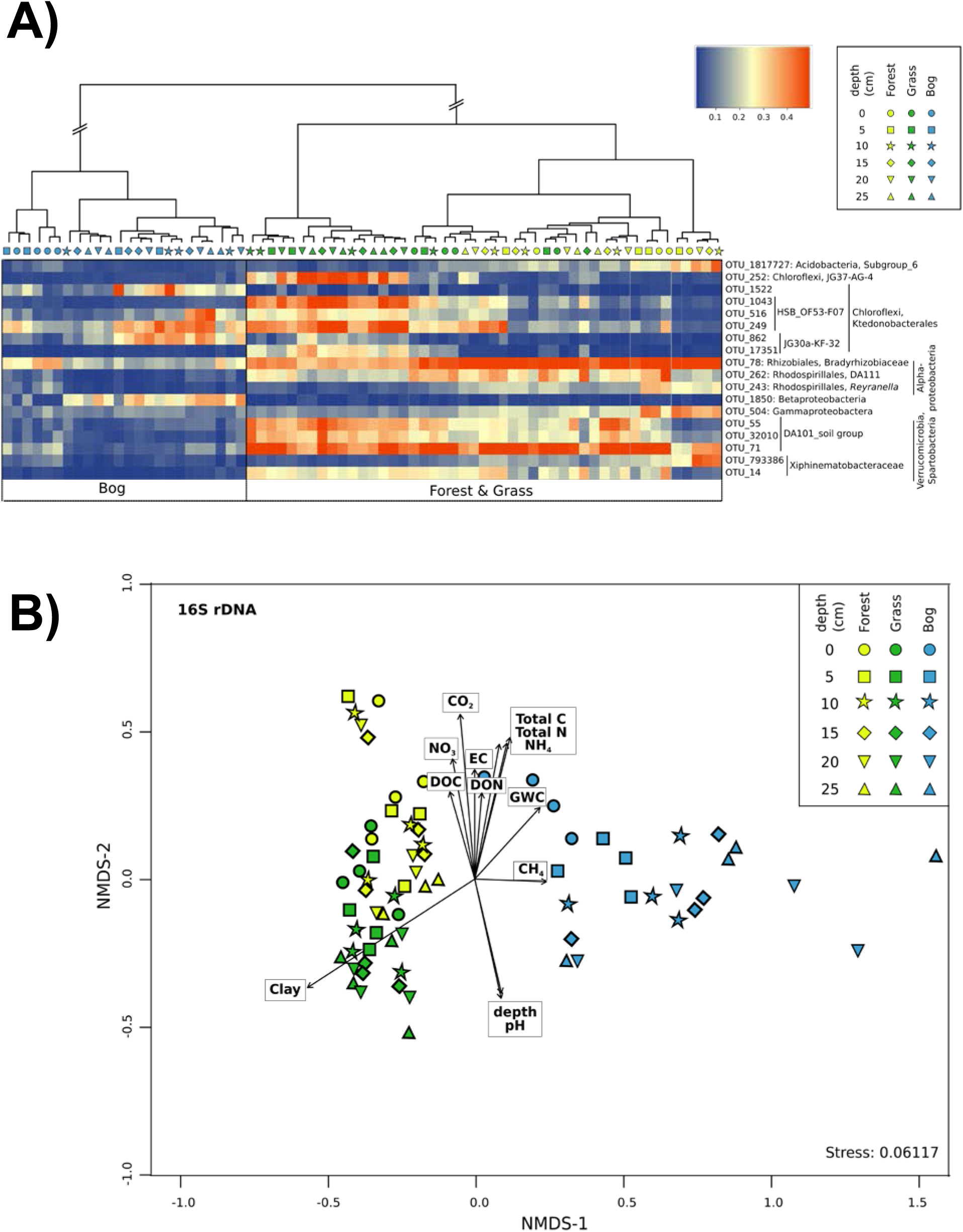
**A)** Heatmap of the most relevant OTUs derived from bacterial 16S rRNA genes. The samples and OTUs were clustered according to Euclidean distances between all Hellinger transformed data. The taxonomy of OTUs was determined using the Sina classifier. The colored scale gives the percentage abundance of OTUs. **B)** NMDS ordination of bacterial 16S rRNA communities based on the Bray–Curtis dissimilarity of community composition (stress =0.061). Arrows indicate the direction at which the environmental vectors fit the best (using the *envfit* function) onto the NMDS ordination space. EC, electrical conductivity; DOC, dissolved organic carbon; DON, dissolved organic nitrogen; GWC, gravimetric water content.

This study focused on methanotroph identification by using Illumina sequencing, of either 16S rRNA or *pmoA* genes. Both 16S rRNA and *pmoA* gene sequence data revealed that the forest and grassland soils were dominated by USCα (Figs. 7, S2) - with nearly equivalent abundances between the two soil types. The highest USCα relative abundance approached 1% of all 16S rRNA genes, and decreased with depth. *Methylocystis* were the most abundant aerobic methanotrophs detected in the bog accounting for up to 0.3% relative abundance across all depths (Fig. 7). *Candidatus* Methylomirabilis sp., which are nitrite-dependent anaerobic methanotrophs, increased with depth in the bog to a maximum of 1.5% relative abundance at 25-30 cm depths. The Illumina sequencing of the *pmoA* gene resulted in distinct differences among all three soil types, but especially between the forest/grassland and bog soils. Using *pmoA*, USCα dominated methanotroph populations in forest and grassland soils, whereas *Methylocystis* dominated bog soils (Fig. S2). Greater counts of USCα were mostly found at the surface of just one bog site (B4). This bog site has greater slope along the soil-vegetation gradient and 80-1200 % greater rock content than the other bog sites across depths (B4; Table S3, Fig 1), thus greater sand content and porosity possibly provided better habitat for methanotrophs in this bog soil.

**Fig. 7.**
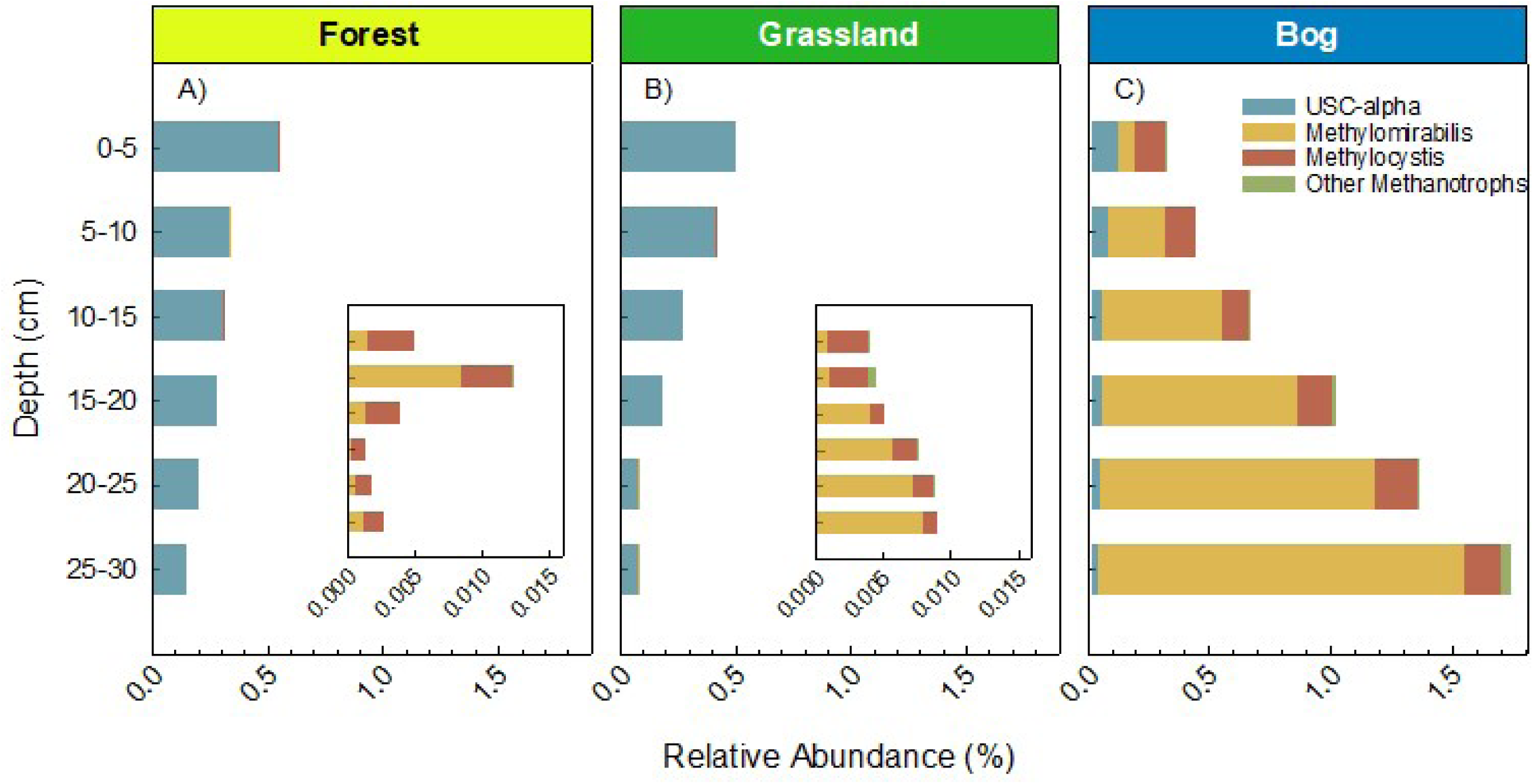
Relative abundance of 16S rRNA genes of the dominant methanotroph groups detected in the forest (**A**), grass (**B**), and bog (**C**) sites. USCα was identified by blast as described in the methods. *Methylocystis* and *Methylomirabilis* were identified based on the Silva classifications. Other methanotrophs include *Methylomonas* and *Methylospira*.

### Patterns in Community Composition and Relationships with Soil Properties

Non-metric dimensional scaling (NMDS) shows a distinct difference in bacterial 16S rRNA genes among soil types and relationship to soil properties (P<0.05, Fig. 6B). The communities in all soils showed a pattern with depth, but the bog soil had the most distinct separation among bacterial communities with depth. Elevated CH_4_ production also correlated with depth of bog soils (Fig. 6B). The grassland soils were most positively related to clay content of the soils, and negatively with soil moisture – these were the driest of the three soil types (Fig. 6B, Table S3). The variables arguably most related to organic matter content (dissolved organic and total C and N, inorganic N, and CO_2_ production) correlated positively with the forest soils, and the somewhat shallow bog soil depths.

The forest *pmoA* community was relatively tightly clustered except for one sample at 10-15 cm depth (Fig. S3A). GHG production did not correlate significantly with the *pmoA* community, indicating differences in the microbial community are not the direct cause of production at depth. The community composition of the Euryarchaeota showed differences among soil types, especially between grassland and bog soils, but no trend with depth (Fig. S3B). Both GHGs and soil properties were positively related to soil organic matter, whereas clay was negatively related to the bog Euryarchaeota community composition. In contrast, the community composition in the grass was positively associated with soil clay content. (Fig. S3B)

## Discussion

### Soil Microbial Community Composition and Links to Greenhouse Gas Fluxes

Adjacent soils, with varying soil-forming factors (especially organisms – vegetation, and landscape position) showed distinct microbial community composition and abundance with depth. The bog soils tended to have the greatest abundance of archaeal 16S rRNA genes (Fig. 4A). At multiple depths, the bog soils also had greater diversity in bacterial and methanotroph communities (Fig. 4B, C). Both of these findings are most likely due to the large total organic C content in bog soils compared to the other two – when averaged across all depths, bog soils had 41 and 128 % greater soil organic C than the forest and grassland soils, respectively (Table S3). The greater the available sources of energy (as approximated by total C) at the base of the food web, the larger and more diverse we might expect the microbial communities to be (Tilman *et al*., 2001; Zhou *et al*., 2002; Hartmann *et al*., 2015).

The bog soils showed the greatest CH_4_ production measured in this study at lower depths (5-10 cm, Fig. 3E), which is surprising considering that at lower depths bog soils have greater water content, i.e., likely more anoxic and methanogenic; however, this can be explained by the increasing abundance of *Candidatus* Methylomirabilis sp. with depth (Fig. 7). *Methylomirabilis* sp. are nitrite-dependent methanotrophs that oxidize CH_4_ by the intracellular production of O_2_ from the dismutation of NO (Ettwig *et al*., 2010). Their presence in this habitat is consistent with previous work that suggests they are common in wetland environments (Hu *et al*., 2014). They occur at deeper depths where nitrite is more available and exogenous O_2_, which is detrimental to their activity (Luesken *et al*., 2012). In addition, methanogenesis might be stimulated at the 5-10 cm depth by Sphagnum rhizoids that occur just above this depth (54%, Table S3). Therefore, high rhizodeposition at this depth could drive methanogenesis by supplying organic C for either acetoclastic or CO_2_ reduction to CH_4_ (Le Mer and Roger, 2001), coupled to relatively low CH_4_ oxidation. Bog soil showed that methanogen gene expression in the top 10 cm of soil correlated linearly with the CH_4_ flux, and methanotroph gene expression ratio was negatively correlated with CH_4_ flux rates at a different peat bog site (Freitag *et al*., 2010). In aggregation, microsites of CH_4_ oxidation and CH_4_ production, when coupled may explain the net production and strong consumption of CH_4_ (Bender and Conrad, 1992, 1994; Von Fischer and Hedin, 2002; Yang and Silver, 2016).

Although not evidence of causation, patterns in soil microbial abundance and community composition were strongly related to GHG production. Given this caveat, the CO_2_ production was best correlated with bacterial 16S rRNA abundance and CH_4_ fluxes with archaeal 16S rRNA abundance (Fig. 5). Archaeal 16S rRNA gene abundance was best related to CH_4_ production (Figs. 3E, 4C, 5E), and highlights the importance of archaeal methanogens in net CH_4_ production (Conrad, 2002; Angel *et al*., 2012; Nazaries, Murrell, *et al*., 2013; Hernández *et al*., 2017).

Methanotroph relative abundance declined with depth in the forest and grassland soil (Fig. 7), similar to the findings from other studies (Bender and Conrad, 1994; Kolb *et al*., 2005). USCα methanotrophs are associated with high rates of atmospheric CH_4_ oxidation in well-drained habitats lacking substantial endogenous methanogenesis (Kolb *et al*., 2005; Chen *et al*., 2007; Bengtson *et al*., 2009; Kolb, 2009; Shrestha *et al*., 2012). The bog soils only showed CH_4_ consumption at the 0-5 cm depth during the summer when soil-to-atmosphere CH_4_ fluxes were highest (Fig. 3A, 3E), which corresponded to the soil depth the greatest proportion of USCα (Fig 7, S2). Thus, USCα abundance, greater O_2_ availability, and lower methanogenesis in the top 0-5 cm all could contribute to the high capacity for aerobic CH_4_ oxidation at the surface of bog soils. Although *Candidatus* Methylomirabilis were highly abundant at depth, their per-cell CH_4_ oxidation capacity might be lower than for aerobic methanotrophs (Luesken *et al*., 2012). Furthermore, the activity of *Candidatus* Methylomirabilis might have been inhibited by O_2_ exposure during the subsectioning of the cores. Finally, although aerobic methanotrophs can survive in low O_2_ environments, in part by energy generation via fermentation reactions (Kalyuzhnaya *et al*., 2013; Kits *et al*., 2015), the rates of CH_4_ oxidation will be lower than most well-oxygenated surface soils.

The three soils had distinct microbial community compositions, with the forest and grassland soils being the most similar (Fig. 6, 7, S2, S3). The bog soils had greater abundances of bacterial 16S rRNA OTU in the family *Ktedonobacterales* (OTU-1522) and an unclassified *Betaproteobacteria* (OTU 1850, Fig. 6A). All soils showed a pattern with depth, but the bog soils showed the greatest discrimination with depth (Fig. 6B). Some microorganisms in the *Ktedonobacterales* have been cultured, and these are Gram-positive, aerobic, broad temperature ranges (meso-to thermophilic), and are characterized by having multicellular filaments and spores – similar to actinomycetes (Yokota, 2012; Yabe *et al*., 2017). The unclassified *Betaproteobacteria* are likely ammonia-oxidizing bacteria and did not play a significant role in CH_4_ oxidation (Bodelier and Frenzel, 1999). The forest soils showed greater abundance of an unclassified *Gammaproteobacteria*, which could potentially be either methanotrophic or ammonia-oxidizing bacteria.

Within the methanotroph community, the bog soils showed a distinctly greater abundance of *Methylocystis* spp., *Ca*. Methylomirabilis, and unclassified proteobacterial methanotrophs (Fig 7 and S2). *Methylocystis* typically proliferate at CH_4_ concentrations > 40ppm, which would occur in bog soils where concentrations (and production) of CH_4_ are greatest. A recent study showed this is indeed the case where low-affinity methanotroph activity is dependent on high supply of CH_4_, and may trigger high-affinity activity during drought (Cai *et al*., 2016). The grassland and forest soils were predominately composed of *Methylocystis* and USCα (Fig 7 and S2). The USCα, are classified as high-affinity, with apparent K_m_ values of 0.01-0.28, compared to that of 0.8-32 for low-affinity methanotrophs (Shukla *et al*., 2013). Several studies now confirm that forest or grassland soils that have high CH_4_ oxidation potential also have high abundance of *Methylocystis* or USCα methanotrophs (Knief *et al*., 2003; Malghani *et al*., 2016). There is mounting evidence, including this study, that indicate the absence/presence of specific methanotrophic bacteria or methanotrophic community composition might be just as important as physicochemical regulators on net CH_4_ fluxes from soils (Nazaries *et al*., 2011; Nazaries, Pan, *et al*., 2013; Malghani *et al*., 2016). Other studies show CH_4_ fluxes are largely regulated by physical processes such as substrate diffusion (Saari *et al*., 1997; Wille *et al*., 2008; Fest *et al*., 2015; D’imperio *et al*., 2017) or labile C supply (Pratscher *et al*., 2011; Sullivan *et al*., 2013); however, due to the high sensitivity of methanotrophic bacteria to physicochemical conditions it is difficult to tease these factors apart (Nazaries *et al*., 2011; Fest *et al*., 2015).

Our study and others (Conrad and Rothfuss, 1991; Frenzel *et al*., 1992) show that surface soils even in consistently wet soils (0-5 cm, Fig. 3) can be a sink not only for atmospheric CH_4_, but also a likely filter for CH_4_ produced at greater depths (Cai *et al*., 2016). Given that by some estimates as much as 80-97% of endogenously produced CH_4_ at depth is consumed before reaching the atmosphere (Conrad and Rothfuss, 1991; Frenzel *et al*., 1992; Oremland and Culbertson, 1992; Sass *et al*., 1992), the disturbance of this thin layer of soil could result in larger net CH_4_ fluxes as CH_4_ produced at depth diffuses toward the atmosphere and is not oxidized in this surface layer. The complex dynamics, high net CH_4_ uptake rates of the forest and grassland soils (Fig. 9A), and high consumption at the surface of the bog soils are all reasons why it is crucial to understand the microbial mechanisms driving these greenhouse gas fluxes.

### Australian Alps: Unique Soils with Disproportionate CH_4_ Sink Strength

Across the globe, aerated upland soils provide a net CH_4_ sink that ranges from 7-100 Tg y^-1^ (Smith *et al*., 2000), which is estimated to be up to 15% of the total global CH_4_ sink (Reeburgh, 2003; Dutaur and Verchot, 2007; Shukla *et al*., 2013). The Australian Alps are restricted to the southeastern corner of the mainland totaling 0.16% of Australia’s area (Fig. 8). We studied 538 ha of a 1.2M ha region, but found convincing evidence for the overwhelming CH_4_ sink strength of this region’s soils (Fig. 9A).

**Fig. 8.**
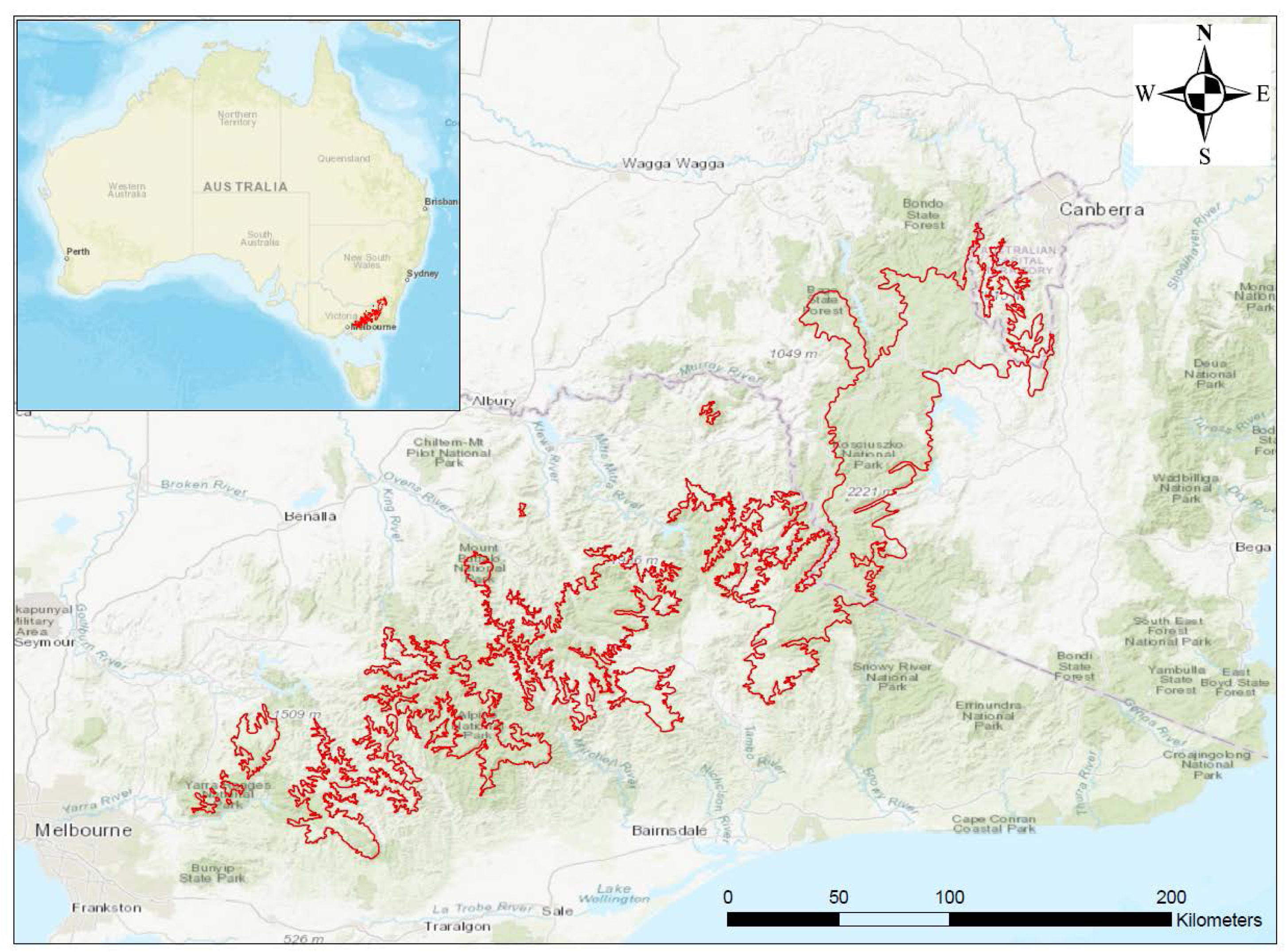
Areal coverage of Australian Alps in southeastern Australia (1.2M ha).

**Fig. 9.**
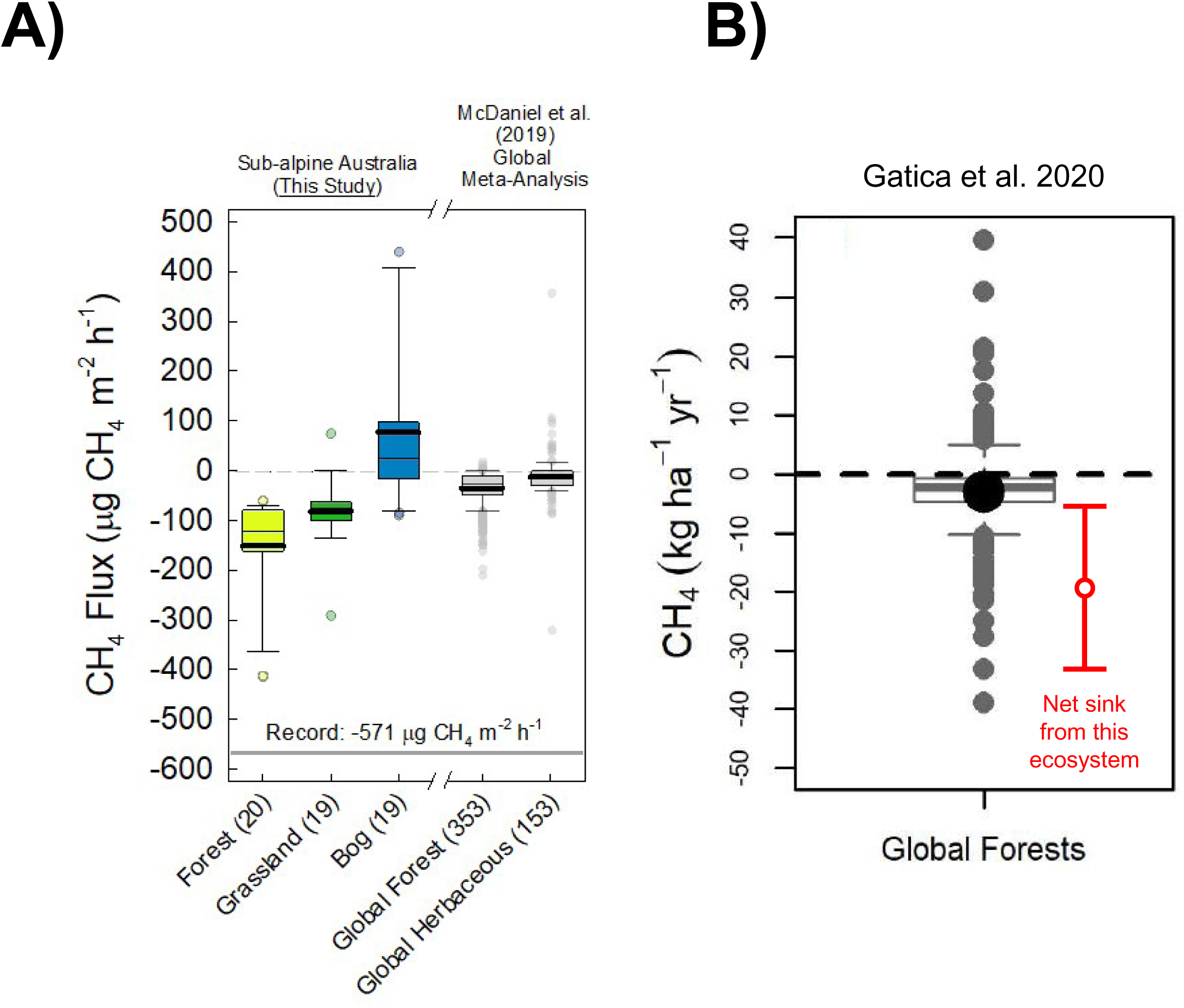
**A)** Hourly methane (CH_4_) fluxes from this study’s Forest, Grassland and Bog soils compared to Forest and Herbaceous studies from a global meta-analysis (McDaniel et al. 2019). 10^th^ and 90^th^ percentile shown by bottom and top whisker. 25^th^ and 75^th^ percentile shown by bottom and top of the box. Median is shown by the thin line, mean by the thick line, and outliers are circles. The number of measurements within each boxplot are shown in parentheses. Gray bar at −571 μg CH_4_ m^-2^ h^-1^ is the greatest CH_4_ oxidation rate (most negative flux) ever observed and published (Singh et al., 1997). **B)** Figure from survey of global forest CH_4_ fluxes (Gatica et al. 2020) with our modeled annual mean net CH_4_ sink from Australian Alps and range created ± relative standard deviation from ecosystem means (Table 2).

We used a combination of foliar coverage estimates for extrapolation across the region, climate station data to interpolate CH_4_ fluxes between seasonal measurements, and range in global sink strength from the literature (Table 2), to parameterize the disproportionate contribution of this endemic ecosystem to the global soil CH_4_ sink. While only comprising <0.03% of global forested and grassland ecosystems, this unique Australian ecosystem could represent 6 up to >100% of global soil methane sink (Kirschke *et al*., 2013; Saunois *et al*., 2016). While it is unreasonable to expect these unique soils to make up more than 100% of the global CH_4_ sink, it illustrates the problem with the low estimates for the global soil CH_4_ sink (Kirschke *et al*., 2013; Saunois *et al*., 2016; Gatica *et al*., 2020). These soils are undoubtedly amongst the greatest negative CH_4_ fluxes observed at an hourly scale (Fig. 9A), and our data conform to other annual CH_4_ flux estimates despite the acknowledged error of these estimates with so few observations (Gatica *et al*., 2020; Fig. 9B). Monitoring CH_4_ fluxes at high spatial and temporal resolution in remote locations remains a major challenge.

Soils under all three vegetation types were sources of CO_2_ with positive GWPs, yet ranged from strong negative to positive GWP contributions from CH_4_ (Table 1). The CH_4_ sink under grassland and forest soils provided 53% and 56% GWP offset (in negative CO_2_-equivalents) of the net CO_2_ emissions respectively. All soils consumed CH_4_ at 0-5 cm depth. This is consistent with other studies showing high uptake in surface soils where O_2_ is more available (Koschorreck and Conrad, 1993; Bender and Conrad, 1994; Hütsch, 1998), even lake sediments can show similar CH_4_ oxidation profiles with depth (He *et al*., 2012). However, many studies find maximum oxidation rates are not near the soil surface, but 5-10 cm below (Bender and Conrad, 1994; Hütsch *et al*., 1994; Schnell and King, 1994; Priemé and Christensen, 1997; Wolf *et al*., 2012). Differences in where this maximum CH_4_ consumption depth occurring are dependent on soil texture, organic matter content, and moisture content with depth along with substrates that are also thought to regulate CH_4_ oxidation like labile C or inorganic N that are also more available in surface soils (Priemé and Christensen, 1997; Pratscher *et al*., 2011; Sullivan *et al*., 2013).

This endemic ecosystem, with an arguably disproportionate importance to the global soil CH_4_ sink (Tables 1,2; Fig. 8), are amongst the ecosystems most likely to be affected by changes in climate, as was recently demonstrated by declines in CH_4_ uptake in several long-term studies of forest soils (Ni and Groffman, 2018). More frequent and intense forest fires are just one global change threat to these Australian Alps soils (Adams, 2013; Adams *et al*., 2013), with potential positive feedbacks to climate change. We are increasingly reliant on key soil ecosystem services to thrive on planet Earth (Jónsson and Davíðsdóttir, 2016), and agroecosystem soils get much of the attention due to proximal benefits (i.e. crop production). Global soil ecosystem services, however, like that of strong atmospheric CH_4_ oxidation occurring in these Australian Alps soils, are not as obvious nor as easy to value but critical for buffering against anthropogenic climate change.

## Experimental Procedures

### Study Site Characteristics

Our study sites were located in an area known locally as the Snowy Plains, within the Snowy Mountains region of southern New South Wales. The Snowy Plains are form part of the Australian Alps montane grasslands, and lie adjacent to the Kosciuszko National Park (36°10ʹ S, 148°54 ʹE; Fig. 1). The elevation range of the sampling area within the Snowy Plains was 1471 to 1677 masl. The mean annual temperature and precipitation are 6.4 °C and ∼1600 mm (SILO - Queensland Government). The site remains snow-covered typically for 2-3 months of the year.

The area is a mosaic ecosystem containing a mixture of bog, grassland, and forest (Fig. 1) and has been well described elsewhere (e.g. Jenkins and Adams, 2010). Alpine humus soils (Chernic tenosol) in the region are ∼400 million years old and derived from glacial moraines of Silurian Mowomba granodiorite (Costin *et al*., 1952). They show little horizon development in the top 30 cm (Fig. S4). Soil are mostly sandy loams in texture, and pH ranges from ∼5.3-5.7 down to 30 cm, across all sites (Table S3). Bogs are dominated by *Sphagnum spp.* and some sparse grass cover (*Poa spp*). Grasslands are dominated mostly by *Poa hiemata* and *Poa costiniana*. Forests are dominated by Snow Gums (*Eucalyptus pauciflora*), with the N-fixing shrub *Bossiaea foliosa*, and some *Poa* spp. as understory and ground layers.

### Experimental Design, Soil Sampling, and Gas Sampling

We used four topographic transects (from bogs and grasslands in the lowest part of the landscape, to forests in upland areas) to guide our soil and GHG sampling (Fig. 1). After each sampling location was determined, a 15-cm diameter (5-cm deep) PVC collar was installed for soil GHG sampling a minimum of 1 month prior to sampling, in order to preclude artifacts introduced by recent disturbance. We measured greenhouse gas fluxes *in situ* and collected soils on 17 February, 25 May, 22 September, and 23 November in 2015. But in-depth microbial sampling and analyses were conducted only on soils collected on 17 February – the time of year for peak CH_4_ production and consumption.

GHG fluxes were measured *in situ* using the static chamber method. We first placed a 3.2 L, vented, PVC chamber over the collar. Four gas samples were collected every 10-15 minutes and directly injected into a Labco Exetainer vial. Concurrent measurements of soil moisture (Theta Probe, Delta-T DevicesTheta Probe, Cambridge, UK) and temperature (Novel Ways, Hamilton, NZ) were taken in triplicate and averaged for each measurement location (7 cm depth).

Immediately after completing gas sampling, the chambers were removed, and a 5-cm diameter 30-cm deep soil core was taken directly in the center of the collar. These cores were used for determination of soil properties, microbial community analyses, and to measure net CO_2_ production and CH_4_ production/consumption in the laboratory. The 30-cm soil cores were stratified and dissected in the field with a clean knife at 5-cm intervals. At each depth, a subsample (∼ 3-5 g of soil) was immediately placed into a 2-ml Eppendorf tube, then placed in liquid nitrogen and stored at −80 °C until DNA was extracted. The remainder of the soil core was placed in a protective PVC sleeve, and then stored in an iced cooler until reaching the laboratory.

Soil cores from the field were transferred to 1 L jars where they were briefly incubated at near-ambient air temperature at the time of collection (22 °C) in order to further characterize CO_2_ and CH_4_ production and consumption occurring at each depth (*sensu* Bender and Conrad, 1994). This procedure has been found to be highly related (r^2^ = 0.44, n = 30) to *in situ* CH_4_ production/consumption (Priemé and Christensen, 1997). The incubations began by placing the relatively undisturbed soil cores into the jars, flushing them with ambient air, and then placing the lids on the jars. The lids had Luer-lock access, and four gas samples were taken over the short-term (3-day) incubations and placed directly into Labco Exetainer vials. Once the incubations finished, the soils were air dried for physical and chemical analyses. The GHG concentrations were analyzed on a gas chromatograph.

### Soil Processing and Analyses

Soils were sieved (2 mm) and both rocks and roots were removed and weighed. A sub-sample was ground for total C and N analysis on a TruSpec Elemental Analyzer (LECO, St. Joseph, MI, USA). Soil texture was analyzed by using the hydrometer method (Gee and Bauder, 1986). Soil inorganic N (ammonium and nitrate) was measured by extracting 5 g of soil with 40 ml of 0.5 M K2SO4 shaken for 1 h and filtered with Whatman #1. These extracts were analyzed on a Lachat Injection-flow Analyzer according to standard methods (Lachat Instruments, Loveland, CO, USA). Dissolved organic C and N were analyzed on the same extracts on a Shimadzu TOC-N Analyzer (Shimadzu Scientific Instruments Inc., Columbia, MD, USA). Electrical conductivity and pH were measured on SevenMulti probe (MettlerToledo, Columbus, OH, USA) with a 1:1 (w:w) ratio with de-ionized water. Heavy elements were analyzed by X-ray fluorescence using a Niton XL3 t Ultra Analyzer meter (Thermo Scientific, Waltham, MA, USA).

### DNA Extraction and Quantitative PCR

Total DNA was extracted using the NucleoSpin^®^ Soil kit (Macherey-Nagel, Germany) by disrupting the cells by bead beating (30 s at 5.5 m s^-1^). DNA purity and quantification were determined using a NanoDrop® Spectrophotomoter ND-1000 (Thermo Fisher Scientific, USA). DNA at a concentration of 10 ng μl^-1^ was stored at −20 °C for further molecular analysis. The abundance of bacterial- and archaeal-16S rRNA genes as well as *pmoA* genes was performed using an iCycler Instrument (BioRad). For all assays, standards containing known number of DNA copies of the target gene were used. qPCR conditions for archaeal- and bacterial-16S rRNA genes were based on dual-labeled probes. For bacterial 16S rRNA genes, primers Bac338F, Bac805R and Bac516P were used. For archaeal 16S rRNA genes, primers Arc 787F, Arc1059R and Arc915P were used. Conditions for both genes were as follows: 0.5 μM of each primer, 0.2 μM of the dual-labeled probe, 3 µl of template, 4 mM MgCl2 (Sigma) and 12.5 µl of JumpStart Ready Mix (Sigma-Aldrich). 1 µl of BSA (0.8 µg/µl) was added to archaeal 16S rRNA gene reactions. The program used for both assays was 94 °C for 5 min, 35 cycles of 95 °C for 30 s and 62 °C for 60 s extension and signal reading (Yu *et al*., 2005). qPCR condition for *pmoA* genes was using SYBR Green (Sigma-Aldrich) and primers A189F / mb661R. PCR conditions were 0.667 µM of each primer, 3 µl template, 4 mM MgCl2 (Sigma), 0.25 µl FITC (1:1000), 12.5 µl of SYBR Green JumpStart *Taq* Ready Mix (Sigma-Aldrich) and 0.6 µl of BSA (0.5 µg/µl). PCR program was 94 °C for 6 min, followed by 45 cycles of 94 °C for 25 s, 65.5 °C 20 s, 72°C 35s, 72°C 10 s plate read. The final melting curve was as follows: 100 cycles of 75-94.8°C 6s, +0.2 °C cycle^-1^ (Kolb *et al*., 2003). Efficiencies of 99.6% for bacterial 16S rRNA genes, 78.8 – 84.9% for archaeal 16S rRNA genes, and 77.2 – 78.5% for *pmoA* genes were obtained, all with R^2^ values > 0.99. Technical duplicates were performed for each of the replicates.

### Illumina library preparation and sequencing

MiSeq Illumina sequencing was performed for total 16S rRNA and *pmoA* genes. PCR primers 515F / 806R targeting the V4 region of the 16S rRNA gene (approximately 250 nucleotides) were used (Bates *et al*., 2011) with an initial denaturation at 94 °C for 5 min, followed by 28 cycles of 94 °C for 30 s, 50 °C for 30 s, and 68 °C for 30 s and a final elongation at 68 °C for 10 min (Hernández *et al*., 2015). The amplification of *pmoA* genes was performed via a semi-nested PCR approach using the primers A189F/A682R for the first round PCR (Holmes *et al*., 1995) as follows: 94 °C for 3 min followed by 30 cycles of 94 °C 45 s, 62 to 52 °C (touchdown 1 °C per cycle) for 60 s, 68 °C 3 min, and a final elongation of 68 °C 10 min (Horz *et al*., 2005). Aliquots of the first round of PCR (0.5 μl) were used as the template in the second round of PCR using the primers A189f / A650r / mb661r in a multiplex PCR as follows: 94 °C for 3 min followed by 25 cycles of 94 °C 45 s, 56 °C 60 s and 68°C 1 min, and final elongation of 68 °C 10 min (Horz *et al*., 2005). Individual PCRs contained a 6-bp molecular barcode integrated in the forward primer. Amplicons were purified using a PCR cleanup kit (Sigma) and quantified using a Qubit 2.0 fluorometer (Invitrogen). An equimolar concentration of the samples was pooled for each of the genes and sequenced on separate runs using 2 x 300 bp MiSeq paired-end protocol. Library preparation and sequencing was performed at the Max Planck Genome Centre (MPGC), Cologne, Germany. Table S4 summarizes primer sequences for both genes and barcode sequences for each of the samples.

### Bioinformatics, Data processing, GIS modeling, and Statistical Analyses

For 16S rRNA genes, quality filtering and trimming forward and reverse adaptors from the sequences was carried out using the tool cutadapt (Martin, 2011). Forward and reverse reads were merged using the usearch fastq_mergepairs command (Edgar, 2013). For the *pmoA* gene, one-end run was performed, and the forward adaptor was trimmed using cutadapt. Downstream processing was performed with UPARSE (Edgar, 2013) and UCHIME pipelines (Edgar *et al*., 2011) following the steps detailed in Reim et al. (Reim *et al*., 2017). For 16S rRNA genes, a representative sequence of each operational taxonomic unit (OTU) was classified based on the SILVA-132 16S rRNA gene database using the naïve Bayesian classifier (bootstrap confidence threshold of 80%) in mothur (Schloss *et al*., 2009). The *pmoA* genes were classified using the same method, but using the *pmoA* database (Dumont *et al*., 2014). Sequence data were deposited in the NCBI Sequence Read Archive (SRA) under accession number PRJNA384296.

A USCα 16S rRNA gene sequence has recently been identified (Pratscher *et al*., 2018), which enabled us to search these sequences in our dataset. The 16S rRNA genes were identified by standalone BLAST against the 16S rRNA OTUs using the USCa_MF sequence (Genbank ID MG203879). Those OTUs with percent ID > 98% relative to the USCa_MF were positively identified as USCα. The relative abundance of USCα in each of the samples could then be calculated from the OTU table.

Greenhouse gas fluxes (both field and incubation) were calculated using a linear regression, or change in the GHG over the time between gas samples. Data was screened for normality and heterogeneity of variances, and when not conforming was log-transformed (Zuur *et al*., 2010) for statistical analyses (all CO2 and gene abundance data). All univariate statistics were conducted in SAS (v. 9.4). Comparisons of variance among soil/vegetation types (bog, grassland, forest) were completed using 1-way ANOVA (α = 0.05) with transect considered random, and depth not included in interactions since depth effects are not independent. Post-hoc tests were completed using lsmeans, adjusted using Tukey’s for multiple comparisons. Multivariate statistics for 16S rRNA Illumina data was analyzed by using the vegan package (Oksanen *et al*., 2016) in R software version 3.0.1 (R Core Team, 2018). Non-metric multidimensional scaling (NMDS) was performed using the *decostand* function for ordination of Hellinger distances. Influence of environmental variables on the total diversity of 16S rRNA and *pmoA* genes was analyzed by the *envfit* function (vegan package in R, permutations = 999). Heatmaps were constructed with the gplots package (Warnes *et al*., 2015). Principal components analysis (PCA) of the Hellinger transformed data was performed using the prcomp function. The OTUs explaining most of the differences between samples were defined as the 10 OTUs contributing the largest absolute loadings in the first and second dimensions of the PCA, obtained from the rotation output file (Hernández *et al*., 2017). Hierarchical clustering of the distance matrix used the “ward.D2” method and *hclust* function. The heatmap was constructed using the *heatmap.2* function in *ggplots* package (Hernández *et al*., 2017).

Using the four field-measured, hourly CH4 fluxes we developed a model to estimate annual CH4 production across the Australian Alps based on number of geographic information system (GIS) datasets. The datasets included elevation and aspect (30 m pixel) as well as daily estimates of maximum and minimum air temperature (BOM, 2020) and soil moisture (0 – 0.1 m; Frost *et al*., 2018). The climate data rasters were all a 5 km pixel size. For each of the days for that CH4 fluxes were determined in the field, the location of 12 sites was used to extract data from each of the raster datasets using ArcGIS (V10.8, ESRI Systems, CA). This data was then used to develop a linear regression model using R (v 4.0.2). The final linear regression model (p-value = <0.0001, F-statistic = 14.95, adjusted r2 = 0.77) included the main factors (the ecosystem measured; Bog, Forest & Grassland) along soil moisture and maximum air temperature and interactions. The spatial extent of each ecosystem across the Australian Alps bioregion (IBRA7, 2012) was based on tree cover in 2015 (DEE, 2018); Forest sites were located in >20% projected foliage cover and Grassland sites <20% projected foliage cover and the Bog sites were considered to be within 1 m of a hydrological flow surface developed using the TauDEM ArcGIS toolset (v5.3.7; Tarboton, 2005). This data along with average seasonal estimates of maximum air temperature and soil moisture were then computed for the Australian Alps. In small number of cases, individual pixels of the maximum air temperature and soil moisture rasters exceeded the maximum/minimum points on which the linear regression model had been developed, in which case they were forced to the maximum/minimum value. The linear regression previously developed was then used to predict seasonal estimate of CH4 fluxes for each of the ecosystems measured across the Australian Alps.

## Acknowledgements

Funding for this research was provided by an anonymous donor. Prof. Dr. Ralf Conrad, at the Max Planck Institute in Marburg, Germany, provided facilities and advice. We would like to kindly thank Barry Aitchison, whom provided us with site access and a wonderful cabin to spend the evenings around the fire. Hero Tahai assisted with laboratory procedures.

## Supplementary Information

**Table S1.**
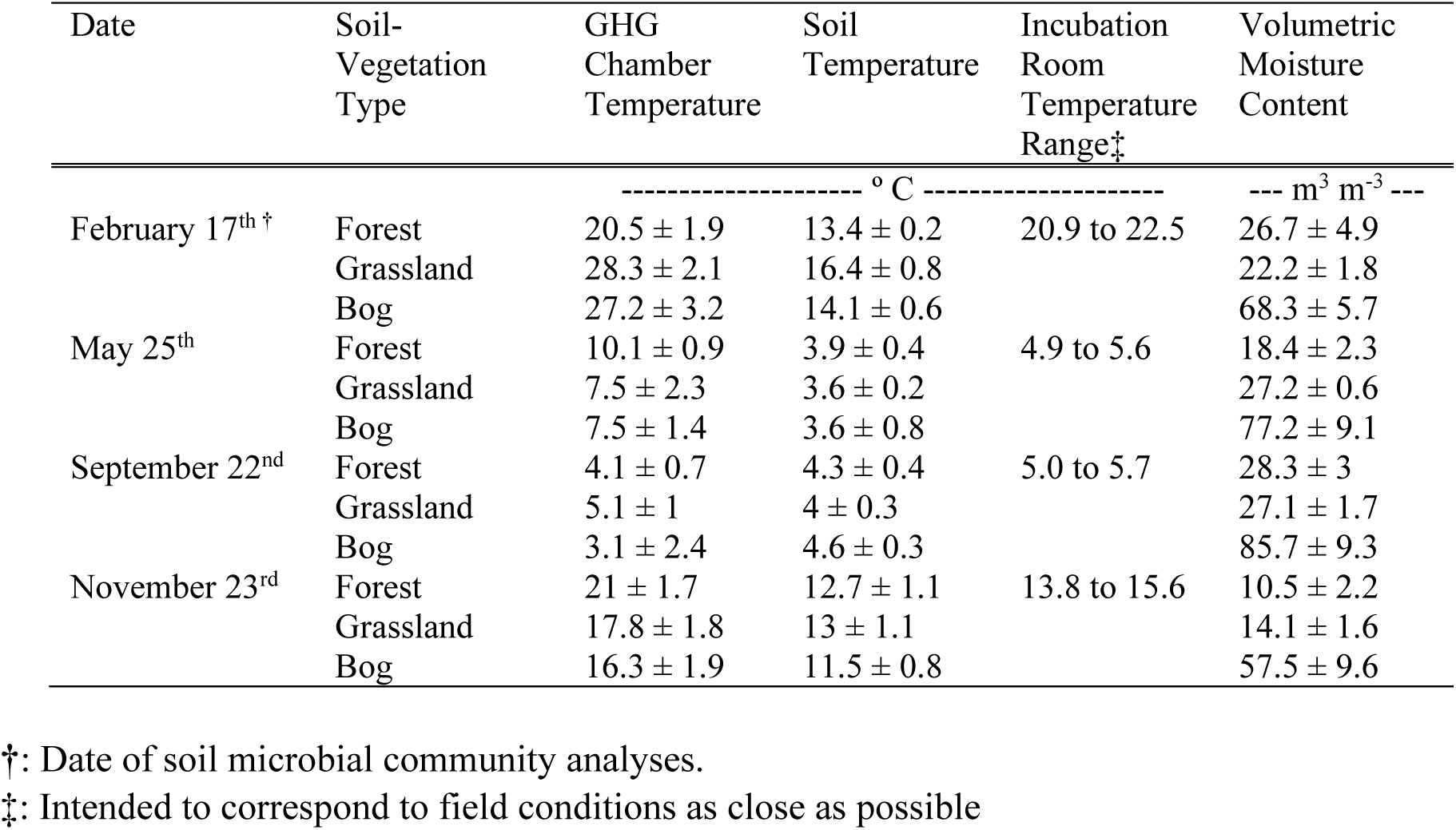
Chamber and soil microclimate at time of sampling, and incubation temperature by date in 2015 (means ± standard errors)^†^

| Date | Soil-Vegetation Type | GHG Chamber Temperature | Soil Temperature | Incubation Room Temperature Range‡ | Volumetric Moisture Content |
| --- | --- | --- | --- | --- | --- |
|  |  | ----- ° C ----- |  |  | --- m<sup>3</sup> m<sup>-3</sup> --- |
| February 17<sup>th</sup> † | Forest | 20.5 ± 1.9 | 13.4 ± 0.2 | 20.9 to 22.5 | 26.7 ± 4.9 |
| | Grassland | 28.3 ± 2.1 | 16.4 ± 0.8 | | 22.2 ± 1.8 |
| | Bog | 27.2 ± 3.2 | 14.1 ± 0.6 | | 68.3 ± 5.7 |
| May 25<sup>th</sup> | Forest | 10.1 ± 0.9 | 3.9 ± 0.4 | 4.9 to 5.6 | 18.4 ± 2.3 |
| | Grassland | 7.5 ± 2.3 | 3.6 ± 0.2 | | 27.2 ± 0.6 |
| | Bog | 7.5 ± 1.4 | 3.6 ± 0.8 | | 77.2 ± 9.1 |
| September 22<sup>nd</sup> | Forest | 4.1 ± 0.7 | 4.3 ± 0.4 | 5.0 to 5.7 | 28.3 ± 3 |
| | Grassland | 5.1 ± 1 | 4 ± 0.3 | | 27.1 ± 1.7 |
| | Bog | 3.1 ± 2.4 | 4.6 ± 0.3 | | 85.7 ± 9.3 |
| November 23<sup>rd</sup> | Forest | 21 ± 1.7 | 12.7 ± 1.1 | 13.8 to 15.6 | 10.5 ± 2.2 |
| | Grassland | 17.8 ± 1.8 | 13 ± 1.1 | | 14.1 ± 1.6 |
| | Bog | 16.3 ± 1.9 | 11.5 ± 0.8 | | 57.5 ± 9.6 |

†: Date of soil microbial community analyses.
‡: Intended to correspond to field conditions as close as possible

**Table S2.**
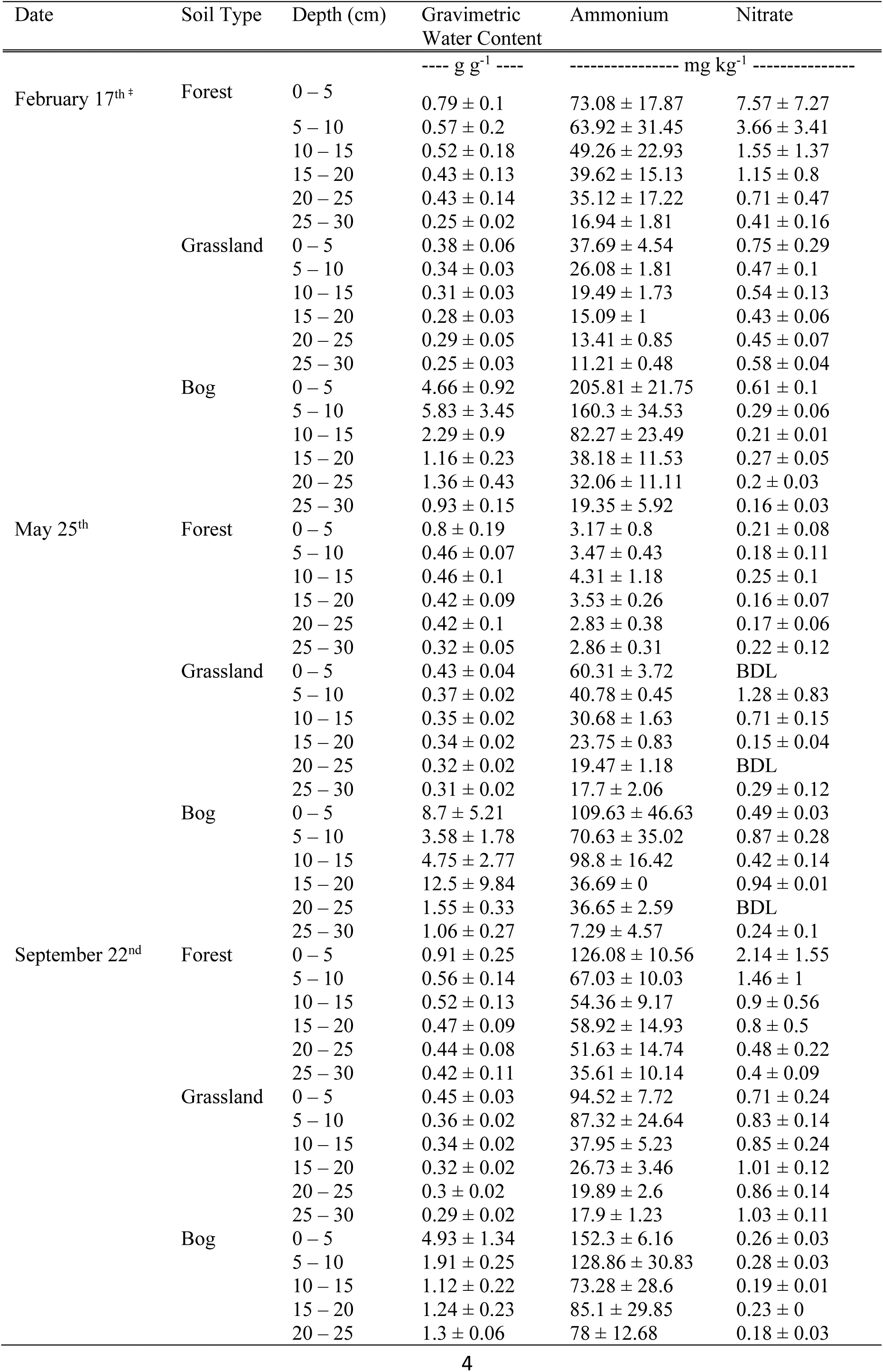
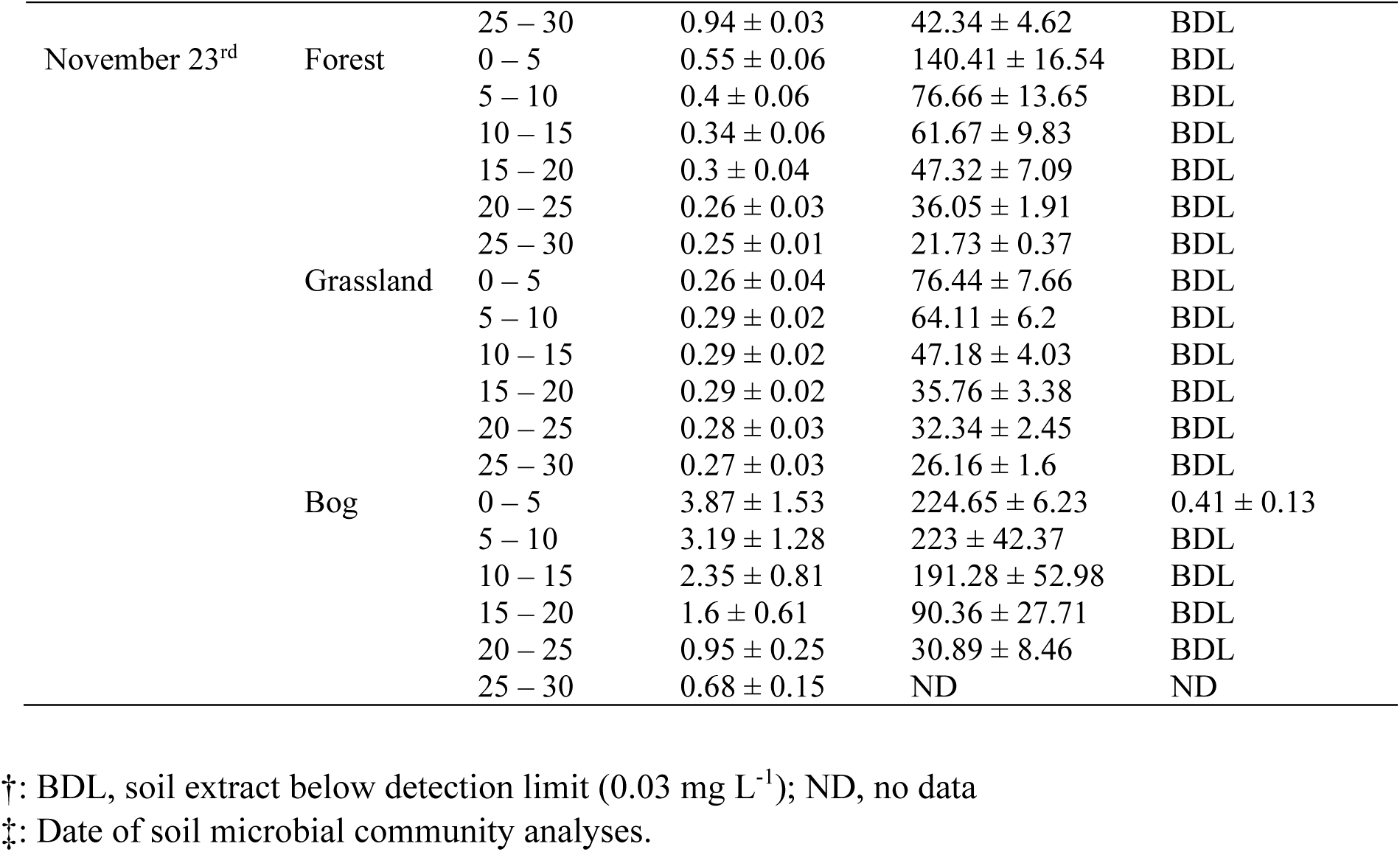
Dynamic soil physical and chemical characteristics by date in 2015 (means ± standard errors)^†^

**Table S3.**
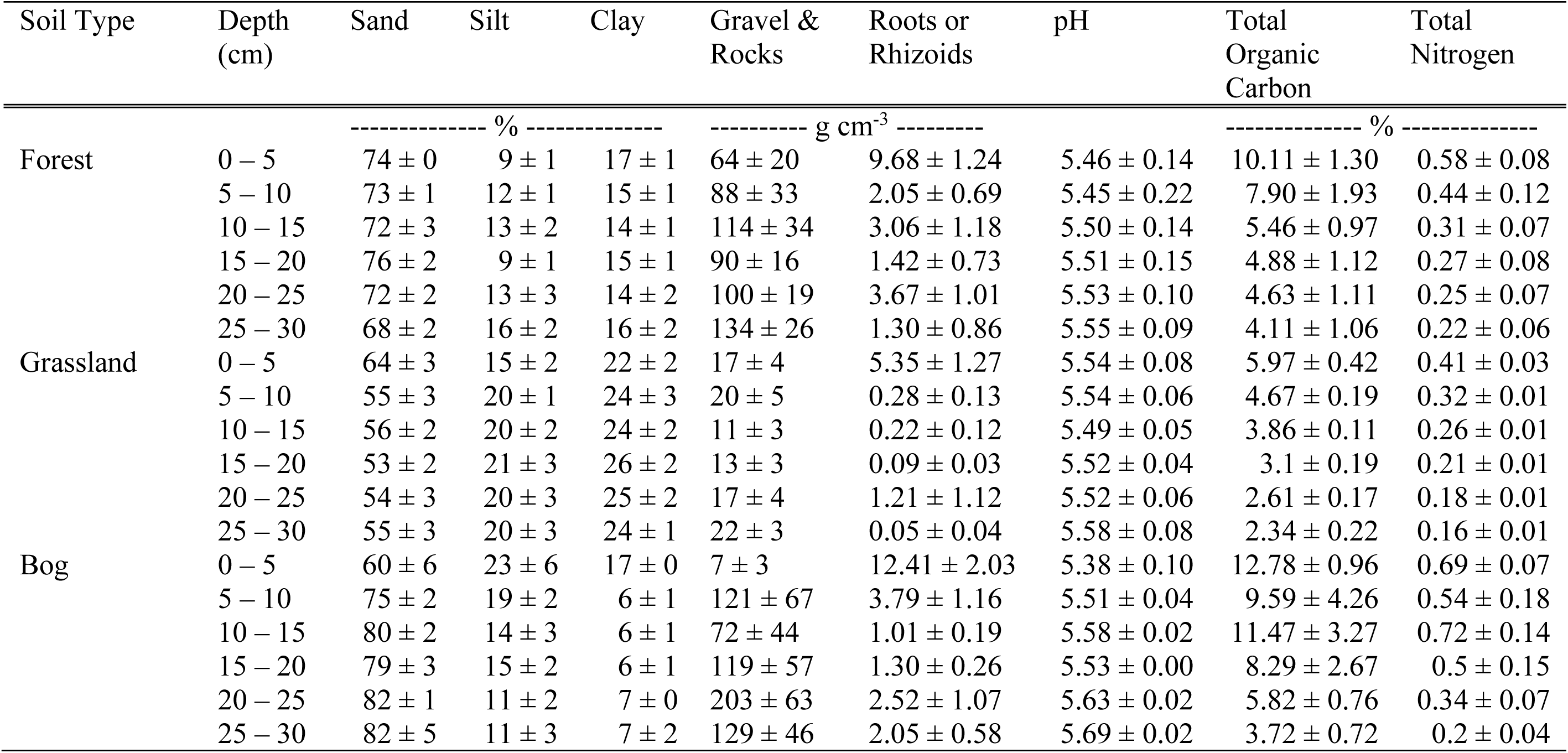
Static soil physical and chemical characteristics – from 17 February 2015 (means ± standard errors)

| Soil Type | Depth<br>(cm) | Sand | Silt | Clay | Gravel &<br>Rocks | Roots or<br>Rhizoids | pH | Total<br>Organic<br>Carbon | Total<br>Nitrogen |
| --- | --- | --- | --- | --- | --- | --- | --- | --- | --- |
|  |  | ----- % ----- |  |  | ----- g cm<sup>-3</sup> ----- |  |  | ----- % ----- |  |
| Forest | 0 – 5 | 74 ± 0 | 9 ± 1 | 17 ± 1 | 64 ± 20 | 9.68 ± 1.24 | 5.46 ± 0.14 | 10.11 ± 1.30 | 0.58 ± 0.08 |
| | 5 – 10 | 73 ± 1 | 12 ± 1 | 15 ± 1 | 88 ± 33 | 2.05 ± 0.69 | 5.45 ± 0.22 | 7.90 ± 1.93 | 0.44 ± 0.12 |
| | 10 – 15 | 72 ± 3 | 13 ± 2 | 14 ± 1 | 114 ± 34 | 3.06 ± 1.18 | 5.50 ± 0.14 | 5.46 ± 0.97 | 0.31 ± 0.07 |
| | 15 – 20 | 76 ± 2 | 9 ± 1 | 15 ± 1 | 90 ± 16 | 1.42 ± 0.73 | 5.51 ± 0.15 | 4.88 ± 1.12 | 0.27 ± 0.08 |
| | 20 – 25 | 72 ± 2 | 13 ± 3 | 14 ± 2 | 100 ± 19 | 3.67 ± 1.01 | 5.53 ± 0.10 | 4.63 ± 1.11 | 0.25 ± 0.07 |
| | 25 – 30 | 68 ± 2 | 16 ± 2 | 16 ± 2 | 134 ± 26 | 1.30 ± 0.86 | 5.55 ± 0.09 | 4.11 ± 1.06 | 0.22 ± 0.06 |
| Grassland | 0 – 5 | 64 ± 3 | 15 ± 2 | 22 ± 2 | 17 ± 4 | 5.35 ± 1.27 | 5.54 ± 0.08 | 5.97 ± 0.42 | 0.41 ± 0.03 |
| | 5 – 10 | 55 ± 3 | 20 ± 1 | 24 ± 3 | 20 ± 5 | 0.28 ± 0.13 | 5.54 ± 0.06 | 4.67 ± 0.19 | 0.32 ± 0.01 |
| | 10 – 15 | 56 ± 2 | 20 ± 2 | 24 ± 2 | 11 ± 3 | 0.22 ± 0.12 | 5.49 ± 0.05 | 3.86 ± 0.11 | 0.26 ± 0.01 |
| | 15 – 20 | 53 ± 2 | 21 ± 3 | 26 ± 2 | 13 ± 3 | 0.09 ± 0.03 | 5.52 ± 0.04 | 3.1 ± 0.19 | 0.21 ± 0.01 |
| | 20 – 25 | 54 ± 3 | 20 ± 3 | 25 ± 2 | 17 ± 4 | 1.21 ± 1.12 | 5.52 ± 0.06 | 2.61 ± 0.17 | 0.18 ± 0.01 |
| | 25 – 30 | 55 ± 3 | 20 ± 3 | 24 ± 1 | 22 ± 3 | 0.05 ± 0.04 | 5.58 ± 0.08 | 2.34 ± 0.22 | 0.16 ± 0.01 |
| Bog | 0 – 5 | 60 ± 6 | 23 ± 6 | 17 ± 0 | 7 ± 3 | 12.41 ± 2.03 | 5.38 ± 0.10 | 12.78 ± 0.96 | 0.69 ± 0.07 |
| | 5 – 10 | 75 ± 2 | 19 ± 2 | 6 ± 1 | 121 ± 67 | 3.79 ± 1.16 | 5.51 ± 0.04 | 9.59 ± 4.26 | 0.54 ± 0.18 |
| | 10 – 15 | 80 ± 2 | 14 ± 3 | 6 ± 1 | 72 ± 44 | 1.01 ± 0.19 | 5.58 ± 0.02 | 11.47 ± 3.27 | 0.72 ± 0.14 |
| | 15 – 20 | 79 ± 3 | 15 ± 2 | 6 ± 1 | 119 ± 57 | 1.30 ± 0.26 | 5.53 ± 0.00 | 8.29 ± 2.67 | 0.5 ± 0.15 |
| | 20 – 25 | 82 ± 1 | 11 ± 2 | 7 ± 0 | 203 ± 63 | 2.52 ± 1.07 | 5.63 ± 0.02 | 5.82 ± 0.76 | 0.34 ± 0.07 |
| | 25 – 30 | 82 ± 5 | 11 ± 3 | 7 ± 2 | 129 ± 46 | 2.05 ± 0.58 | 5.69 ± 0.02 | 3.72 ± 0.72 | 0.2 ± 0.04 |

**Table S4.**
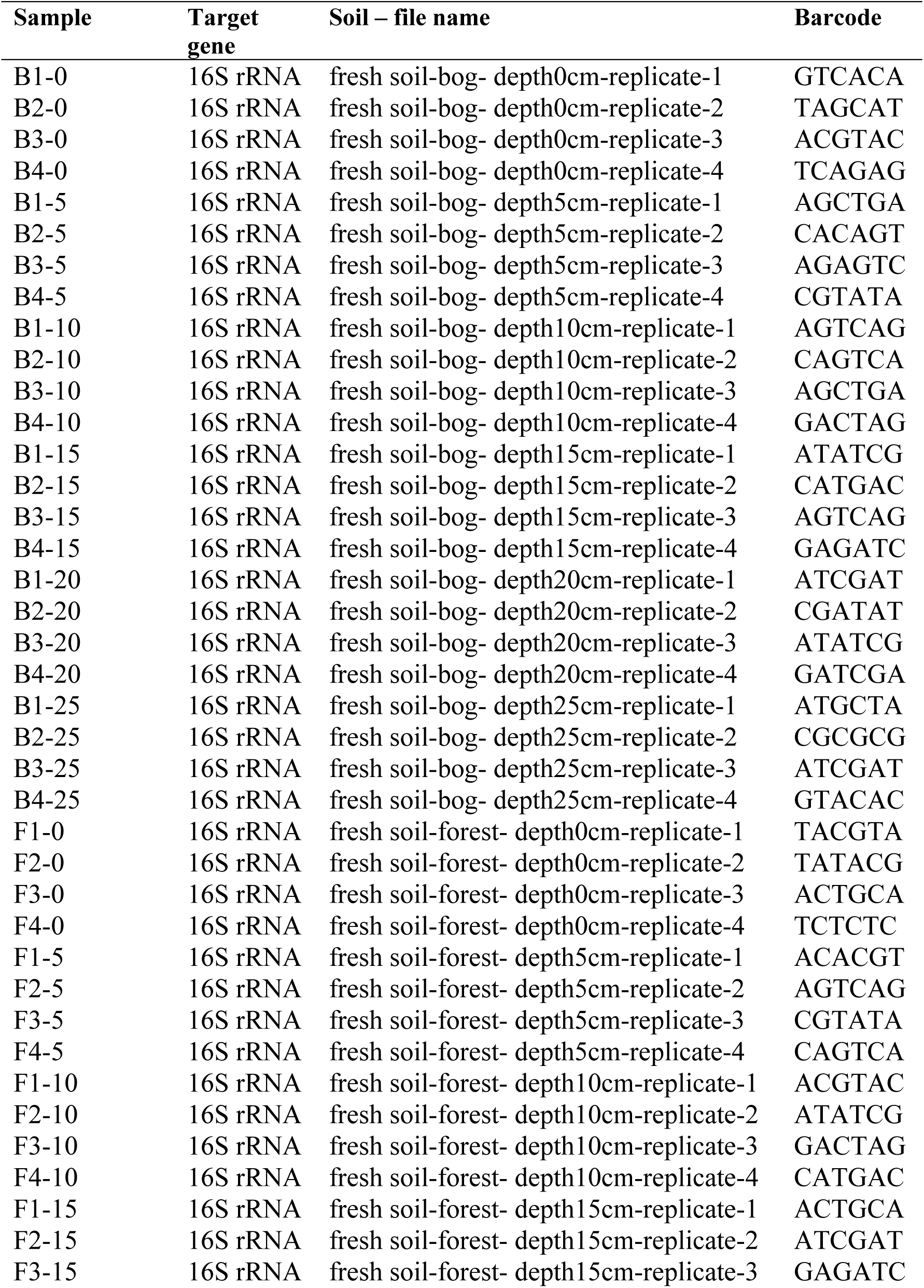
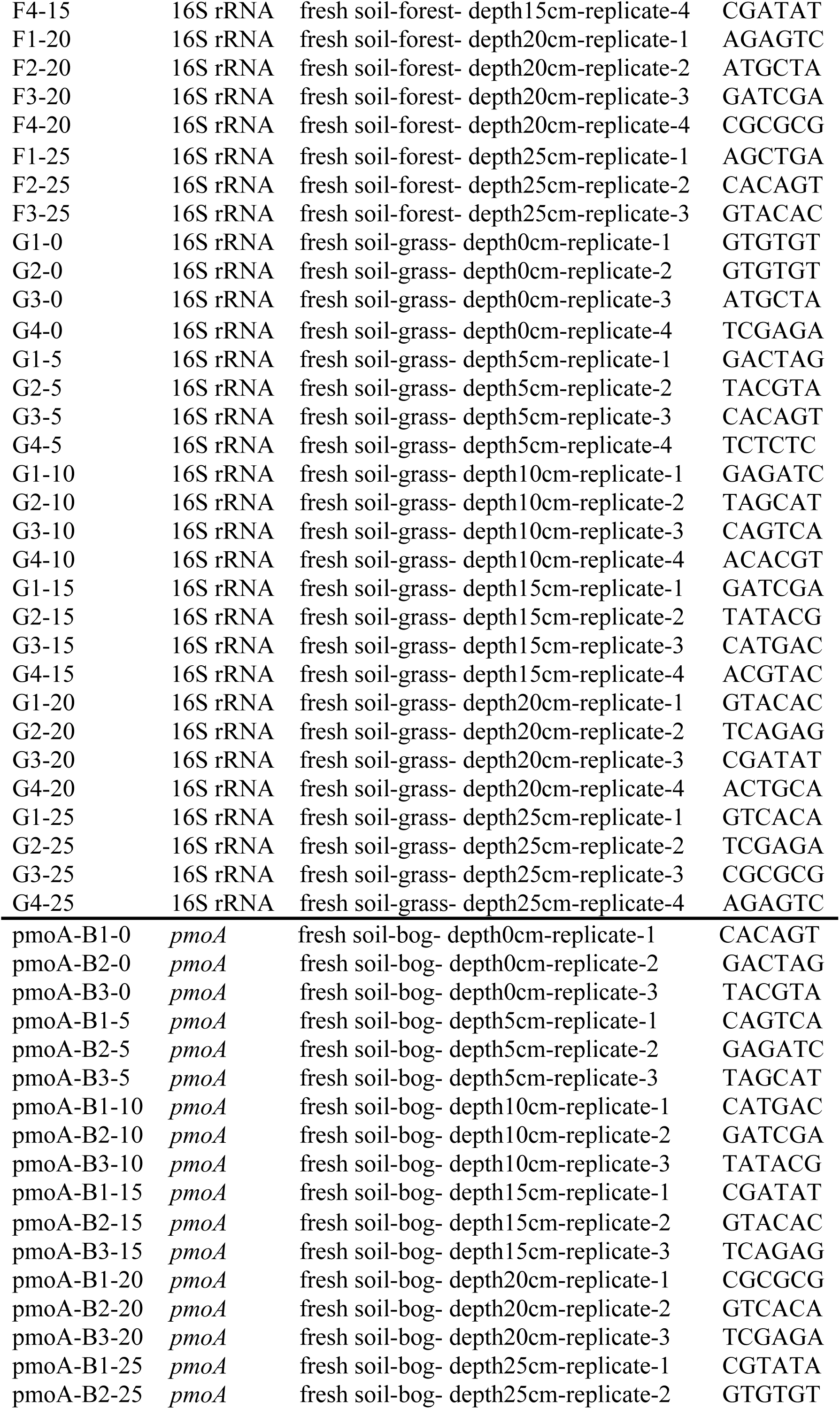
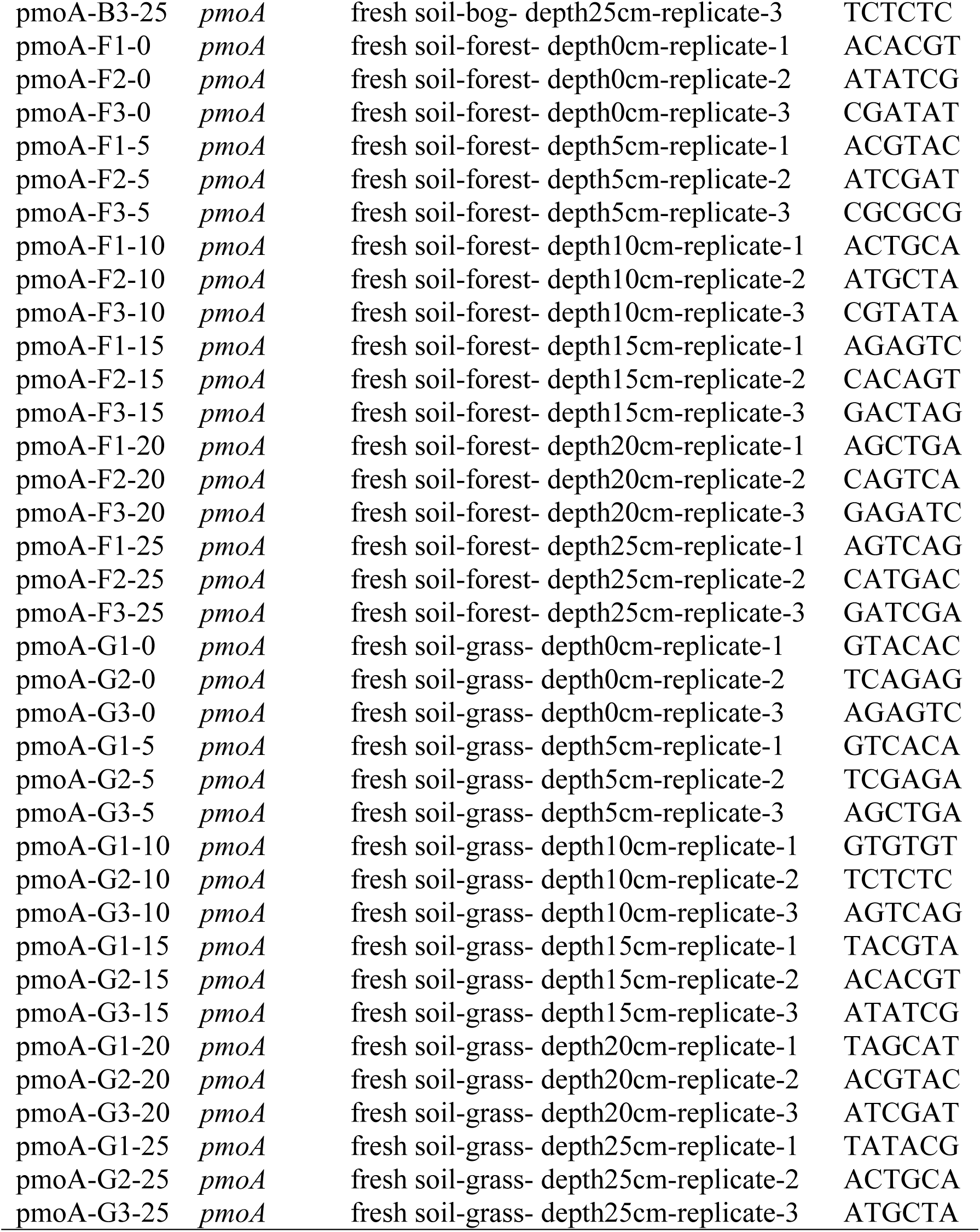
Barcode identification for each of the samples analyzed. Raw data were deposited under the study accession number PRJNA384296 in the NCBI Sequence Read Archive (SRA). For 16S rRNA genes, primers used: F515 (5’-GTGCCAGCMGCCGCGGTAA-3’), R806 (5’-GGACTACVSGGGTATCTAAT-3’). For *pmoA* genes, primer set first round PCR (A189f/A682r) and second round multiplex PCR (A189f/A650r/mb661r): A189f (5’-GGNGACTGGGACTTCTGG-3’), A682r (5’-GAASGCNGAGAAGAASGC-3’), A650r (5’-ACGTCCTTACCGAAGGT-3’), mb661r (5’-CCGGMGCAACGTCYTTACC-3’).

**Figure S1.**
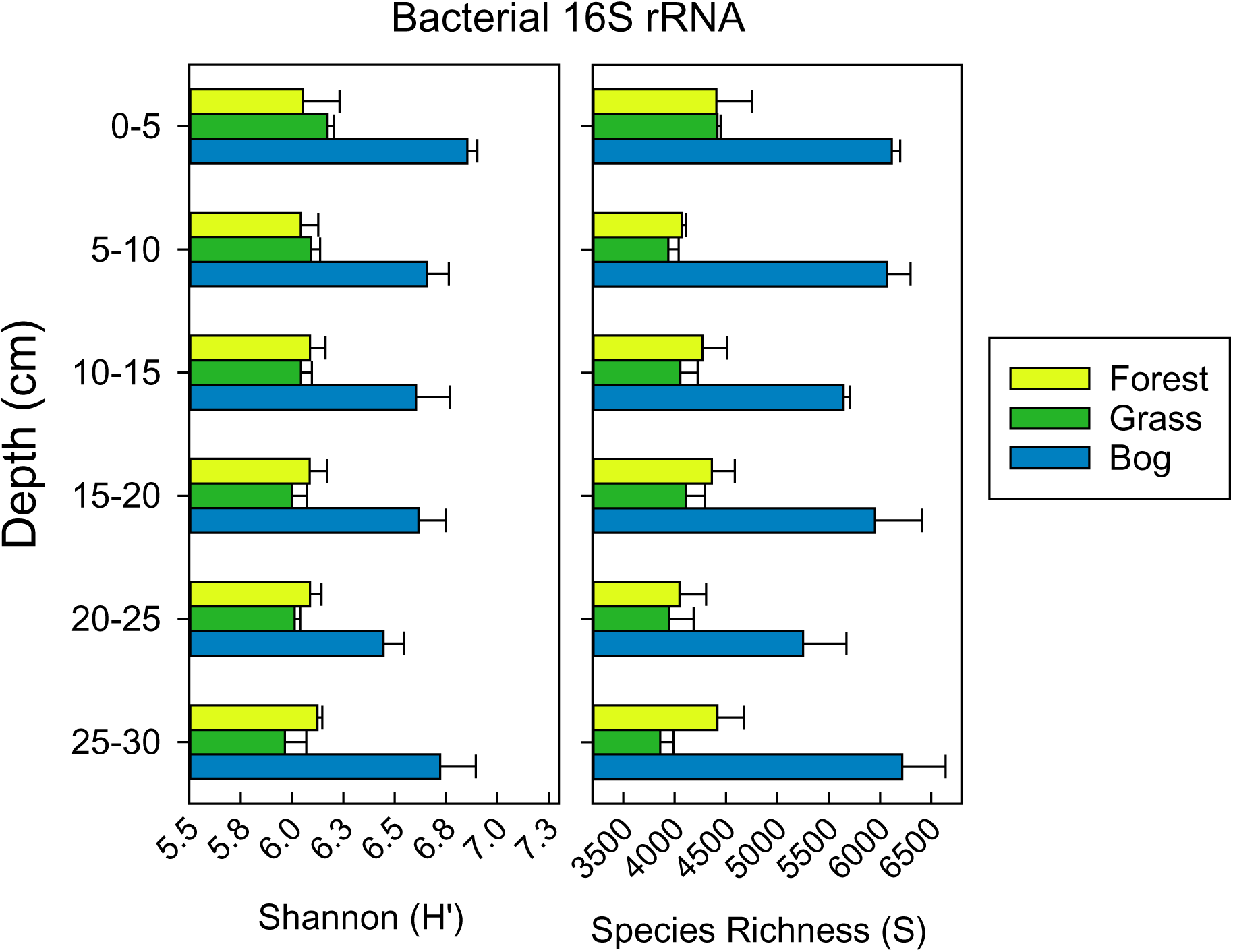
Bacterial 16S rRNA gene diversity measured as Shannon Diversity (H’) and Richness (S) (n=4, means ± standard errors).

**Figure S2.**
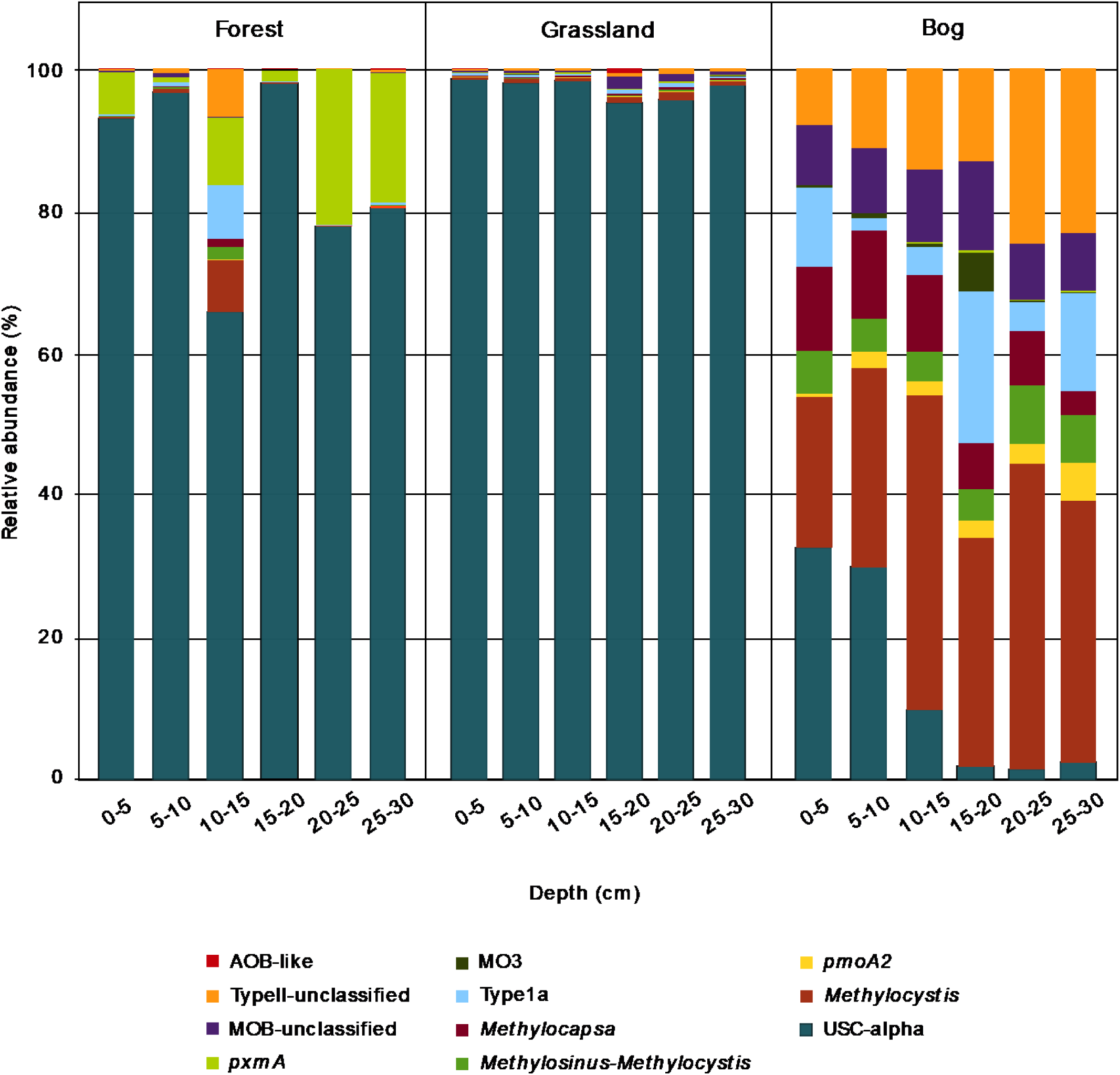
Relative abundance of *pmoA* genes of the dominant methanotroph groups.

**Figure S3.**
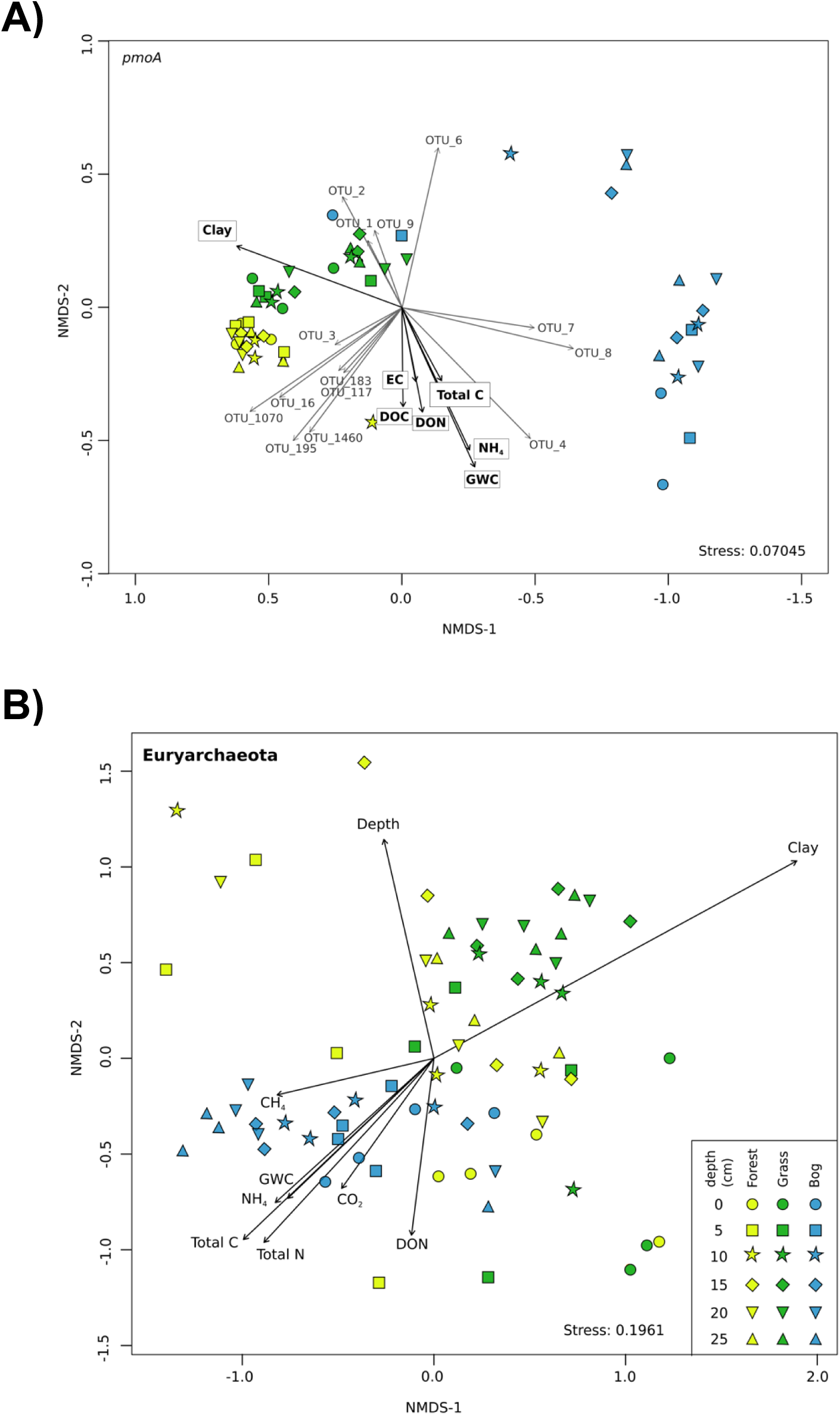
NMDS ordination of *pmoA* (A) and euryarchaeota (B) communities based on the Bray–Curtis dissimilarity of community composition. Shape indicates depth and sites are colored according to the soil typel. The arrows indicate the direction at which the environmental vectors fit the best (using the *envfit* function) onto the NMDS ordination space. Abbreviations: DOC, dissolved organic carbon; DON, dissolved organic nitrogen; EC, electrical conductivity; GWC, gravimetric water content; NH4, ammonium.

**Figure S4.**
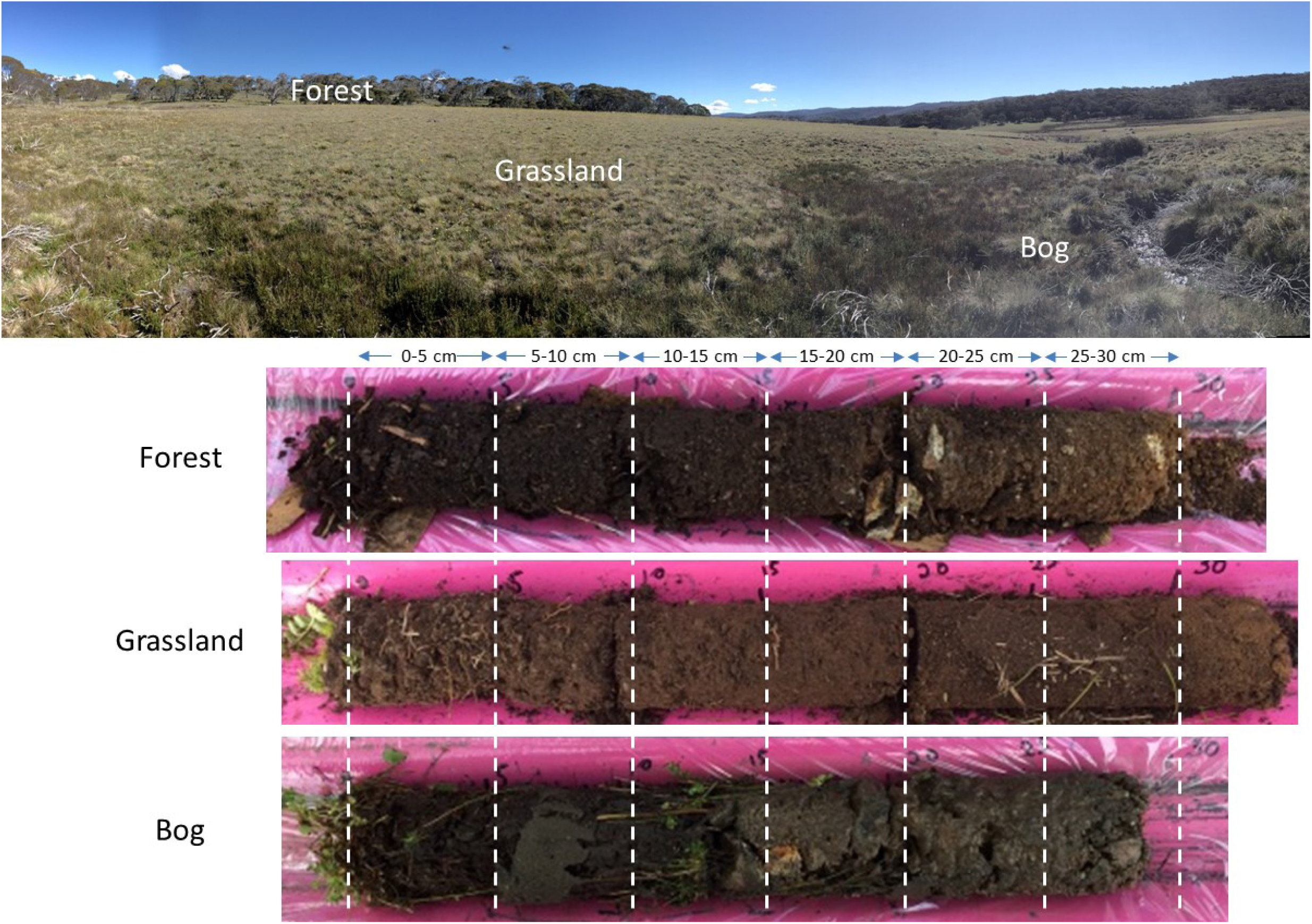
Soil/Vegetation gradient (Forest-Grassland-Bog) located in Snowy Mountains region near Kosciuszko National Park, NSW, Australia. Soil/vegetation types shown above, below are three soil cores from each of the soil/vegetation types.

