## Supplementary for "Disproportionate CH_4_ sink strength from an endemic, sub-alpine Australian soil microbial community"

**Supplementary Information Table of Contents**

### TABLES

**Figure S3**. NMDS ordination of *pmoA* (A) and euryarchaeota (B) communities based on the Bray–Curtis dissimilarity of community composition. Shape indicates depth and sites are colored according to the soil typel. The arrows indicate the direction at which the environmental vectors fit the best (using the *envfit* function) onto the NMDS ordination space. Abbreviations: DOC, dissolved organic carbon; DON, dissolved organic nitrogen; EC, electrical conductivity; GWC, gravimetric water content; $NH_4$, ammonium. ........ 12

**Figure S4.** Example of sampling locations and streams located in a High Country region near Mt. Kosciuszko National Park, NSW, Australia. Soil/vegetation types shown along one transect in the picture above. Below are three soil cores from each of the soil/vegetation types. ........ 13

**Table S1**. Chamber and soil microclimate at time of sampling, and incubation temperature by date in 2015 (means ± standard errors)†

| Date | Soil-Vegetation Type | GHG Chamber Temperature | Soil Temperature | Incubation Room Temperature Range‡ | Volumetric Moisture Content |
|---|---|---|---|---|---|
| | | -------------------- ° C -------------------- | | | --- $m^3$ $m^{-3}$ --- |
| February 17th † | Forest | 20.5 ± 1.9 | 13.4 ± 0.2 | 20.9 to 22.5 | 26.7 ± 4.9 |
| | Grassland | 28.3 ± 2.1 | 16.4 ± 0.8 | | 22.2 ± 1.8 |
| | Bog | 27.2 ± 3.2 | 14.1 ± 0.6 | | 68.3 ± 5.7 |
| May 25th | Forest | 10.1 ± 0.9 | 3.9 ± 0.4 | 4.9 to 5.6 | 18.4 ± 2.3 |
| | Grassland | 7.5 ± 2.3 | 3.6 ± 0.2 | | 27.2 ± 0.6 |
| | Bog | 7.5 ± 1.4 | 3.6 ± 0.8 | | 77.2 ± 9.1 |
| September 22nd | Forest | 4.1 ± 0.7 | 4.3 ± 0.4 | 5.0 to 5.7 | 28.3 ± 3 |
| | Grassland | 5.1 ± 1 | 4 ± 0.3 | | 27.1 ± 1.7 |
| | Bog | 3.1 ± 2.4 | 4.6 ± 0.3 | | 85.7 ± 9.3 |
| November 23rd | Forest | 21 ± 1.7 | 12.7 ± 1.1 | 13.8 to 15.6 | 10.5 ± 2.2 |
| | Grassland | 17.8 ± 1.8 | 13 ± 1.1 | | 14.1 ± 1.6 |
| | Bog | 16.3 ± 1.9 | 11.5 ± 0.8 | | 57.5 ± 9.6 |

†: Date of soil microbial community analyses.
‡: Intended to correspond to field conditions as close as possible

**Table S2**. Dynamic soil physical and chemical characteristics by date in 2015 (means ± standard errors)[†]

| Date | Soil Type | Depth (cm) | Gravimetric Water Content | Ammonium | Nitrate |
|---|---|---|---|---|---|
| | | | ---- $g\ g^{-1}$ ---- | ---------------- $mg\ kg^{-1}$ --------------- | |
| February 17th [‡] | Forest | 0 – 5 | 0.79 ± 0.1 | 73.08 ± 17.87 | 7.57 ± 7.27 |
| | | 5 – 10 | 0.57 ± 0.2 | 63.92 ± 31.45 | 3.66 ± 3.41 |
| | | 10 – 15 | 0.52 ± 0.18 | 49.26 ± 22.93 | 1.55 ± 1.37 |
| | | 15 – 20 | 0.43 ± 0.13 | 39.62 ± 15.13 | 1.15 ± 0.8 |
| | | 20 – 25 | 0.43 ± 0.14 | 35.12 ± 17.22 | 0.71 ± 0.47 |
| | | 25 – 30 | 0.25 ± 0.02 | 16.94 ± 1.81 | 0.41 ± 0.16 |
| | Grassland | 0 – 5 | 0.38 ± 0.06 | 37.69 ± 4.54 | 0.75 ± 0.29 |
| | | 5 – 10 | 0.34 ± 0.03 | 26.08 ± 1.81 | 0.47 ± 0.1 |
| | | 10 – 15 | 0.31 ± 0.03 | 19.49 ± 1.73 | 0.54 ± 0.13 |
| | | 15 – 20 | 0.28 ± 0.03 | 15.09 ± 1 | 0.43 ± 0.06 |
| | | 20 – 25 | 0.29 ± 0.05 | 13.41 ± 0.85 | 0.45 ± 0.07 |
| | | 25 – 30 | 0.25 ± 0.03 | 11.21 ± 0.48 | 0.58 ± 0.04 |
| | Bog | 0 – 5 | 4.66 ± 0.92 | 205.81 ± 21.75 | 0.61 ± 0.1 |
| | | 5 – 10 | 5.83 ± 3.45 | 160.3 ± 34.53 | 0.29 ± 0.06 |
| | | 10 – 15 | 2.29 ± 0.9 | 82.27 ± 23.49 | 0.21 ± 0.01 |
| | | 15 – 20 | 1.16 ± 0.23 | 38.18 ± 11.53 | 0.27 ± 0.05 |
| | | 20 – 25 | 1.36 ± 0.43 | 32.06 ± 11.11 | 0.2 ± 0.03 |
| | | 25 – 30 | 0.93 ± 0.15 | 19.35 ± 5.92 | 0.16 ± 0.03 |
| May 25th | Forest | 0 – 5 | 0.8 ± 0.19 | 3.17 ± 0.8 | 0.21 ± 0.08 |
| | | 5 – 10 | 0.46 ± 0.07 | 3.47 ± 0.43 | 0.18 ± 0.11 |
| | | 10 – 15 | 0.46 ± 0.1 | 4.31 ± 1.18 | 0.25 ± 0.1 |
| | | 15 – 20 | 0.42 ± 0.09 | 3.53 ± 0.26 | 0.16 ± 0.07 |
| | | 20 – 25 | 0.42 ± 0.1 | 2.83 ± 0.38 | 0.17 ± 0.06 |
| | | 25 – 30 | 0.32 ± 0.05 | 2.86 ± 0.31 | 0.22 ± 0.12 |
| | Grassland | 0 – 5 | 0.43 ± 0.04 | 60.31 ± 3.72 | BDL |
| | | 5 – 10 | 0.37 ± 0.02 | 40.78 ± 0.45 | 1.28 ± 0.83 |
| | | 10 – 15 | 0.35 ± 0.02 | 30.68 ± 1.63 | 0.71 ± 0.15 |
| | | 15 – 20 | 0.34 ± 0.02 | 23.75 ± 0.83 | 0.15 ± 0.04 |
| | | 20 – 25 | 0.32 ± 0.02 | 19.47 ± 1.18 | BDL |
| | | 25 – 30 | 0.31 ± 0.02 | 17.7 ± 2.06 | 0.29 ± 0.12 |
| | Bog | 0 – 5 | 8.7 ± 5.21 | 109.63 ± 46.63 | 0.49 ± 0.03 |
| | | 5 – 10 | 3.58 ± 1.78 | 70.63 ± 35.02 | 0.87 ± 0.28 |
| | | 10 – 15 | 4.75 ± 2.77 | 98.8 ± 16.42 | 0.42 ± 0.14 |
| | | 15 – 20 | 12.5 ± 9.84 | 36.69 ± 0 | 0.94 ± 0.01 |
| | | 20 – 25 | 1.55 ± 0.33 | 36.65 ± 2.59 | BDL |
| | | 25 – 30 | 1.06 ± 0.27 | 7.29 ± 4.57 | 0.24 ± 0.1 |
| September 22nd | Forest | 0 – 5 | 0.91 ± 0.25 | 126.08 ± 10.56 | 2.14 ± 1.55 |
| | | 5 – 10 | 0.56 ± 0.14 | 67.03 ± 10.03 | 1.46 ± 1 |
| | | 10 – 15 | 0.52 ± 0.13 | 54.36 ± 9.17 | 0.9 ± 0.56 |
| | | 15 – 20 | 0.47 ± 0.09 | 58.92 ± 14.93 | 0.8 ± 0.5 |
| | | 20 – 25 | 0.44 ± 0.08 | 51.63 ± 14.74 | 0.48 ± 0.22 |
| | | 25 – 30 | 0.42 ± 0.11 | 35.61 ± 10.14 | 0.4 ± 0.09 |
| | Grassland | 0 – 5 | 0.45 ± 0.03 | 94.52 ± 7.72 | 0.71 ± 0.24 |
| | | 5 – 10 | 0.36 ± 0.02 | 87.32 ± 24.64 | 0.83 ± 0.14 |
| | | 10 – 15 | 0.34 ± 0.02 | 37.95 ± 5.23 | 0.85 ± 0.24 |
| | | 15 – 20 | 0.32 ± 0.02 | 26.73 ± 3.46 | 1.01 ± 0.12 |
| | | 20 – 25 | 0.3 ± 0.02 | 19.89 ± 2.6 | 0.86 ± 0.14 |
| | | 25 – 30 | 0.29 ± 0.02 | 17.9 ± 1.23 | 1.03 ± 0.11 |
| | Bog | 0 – 5 | 4.93 ± 1.34 | 152.3 ± 6.16 | 0.26 ± 0.03 |
| | | 5 – 10 | 1.91 ± 0.25 | 128.86 ± 30.83 | 0.28 ± 0.03 |
| | | 10 – 15 | 1.12 ± 0.22 | 73.28 ± 28.6 | 0.19 ± 0.01 |
| | | 15 – 20 | 1.24 ± 0.23 | 85.1 ± 29.85 | 0.23 ± 0 |
| | | 20 – 25 | 1.3 ± 0.06 | 78 ± 12.68 | 0.18 ± 0.03 |

| | | | | | |
|---|---|---|---|---|---|
| | | 25 – 30 | 0.94 ± 0.03 | 42.34 ± 4.62 | BDL |
| November 23rd | Forest | 0 – 5 | 0.55 ± 0.06 | 140.41 ± 16.54 | BDL |
| | | 5 – 10 | 0.4 ± 0.06 | 76.66 ± 13.65 | BDL |
| | | 10 – 15 | 0.34 ± 0.06 | 61.67 ± 9.83 | BDL |
| | | 15 – 20 | 0.3 ± 0.04 | 47.32 ± 7.09 | BDL |
| | | 20 – 25 | 0.26 ± 0.03 | 36.05 ± 1.91 | BDL |
| | | 25 – 30 | 0.25 ± 0.01 | 21.73 ± 0.37 | BDL |
| | Grassland | 0 – 5 | 0.26 ± 0.04 | 76.44 ± 7.66 | BDL |
| | | 5 – 10 | 0.29 ± 0.02 | 64.11 ± 6.2 | BDL |
| | | 10 – 15 | 0.29 ± 0.02 | 47.18 ± 4.03 | BDL |
| | | 15 – 20 | 0.29 ± 0.02 | 35.76 ± 3.38 | BDL |
| | | 20 – 25 | 0.28 ± 0.03 | 32.34 ± 2.45 | BDL |
| | | 25 – 30 | 0.27 ± 0.03 | 26.16 ± 1.6 | BDL |
| | Bog | 0 – 5 | 3.87 ± 1.53 | 224.65 ± 6.23 | 0.41 ± 0.13 |
| | | 5 – 10 | 3.19 ± 1.28 | 223 ± 42.37 | BDL |
| | | 10 – 15 | 2.35 ± 0.81 | 191.28 ± 52.98 | BDL |
| | | 15 – 20 | 1.6 ± 0.61 | 90.36 ± 27.71 | BDL |
| | | 20 – 25 | 0.95 ± 0.25 | 30.89 ± 8.46 | BDL |
| | | 25 – 30 | 0.68 ± 0.15 | ND | ND |

†: BDL, soil extract below detection limit (0.03 mg $L^{-1}$); ND, no data

‡: Date of soil microbial community analyses.

**Table S3**. Static soil physical and chemical characteristics – from 17 February 2015 (means ± standard errors)

| Soil Type | Depth (cm) | Sand | Silt | Clay | Gravel & Rocks | Roots or Rhizoids | pH | Total Organic Carbon | Total Nitrogen |
|---|---|---|---|---|---|---|---|---|---|
| | | -------------- % -------------- | | | ---------- g cm$^{-3}$ --------- | | | -------------- % -------------- | |
| Forest | 0 – 5 | 74 ± 0 | 9 ± 1 | 17 ± 1 | 64 ± 20 | 9.68 ± 1.24 | 5.46 ± 0.14 | 10.11 ± 1.30 | 0.58 ± 0.08 |
| | 5 – 10 | 73 ± 1 | 12 ± 1 | 15 ± 1 | 88 ± 33 | 2.05 ± 0.69 | 5.45 ± 0.22 | 7.90 ± 1.93 | 0.44 ± 0.12 |
| | 10 – 15 | 72 ± 3 | 13 ± 2 | 14 ± 1 | 114 ± 34 | 3.06 ± 1.18 | 5.50 ± 0.14 | 5.46 ± 0.97 | 0.31 ± 0.07 |
| | 15 – 20 | 76 ± 2 | 9 ± 1 | 15 ± 1 | 90 ± 16 | 1.42 ± 0.73 | 5.51 ± 0.15 | 4.88 ± 1.12 | 0.27 ± 0.08 |
| | 20 – 25 | 72 ± 2 | 13 ± 3 | 14 ± 2 | 100 ± 19 | 3.67 ± 1.01 | 5.53 ± 0.10 | 4.63 ± 1.11 | 0.25 ± 0.07 |
| | 25 – 30 | 68 ± 2 | 16 ± 2 | 16 ± 2 | 134 ± 26 | 1.30 ± 0.86 | 5.55 ± 0.09 | 4.11 ± 1.06 | 0.22 ± 0.06 |
| Grassland | 0 – 5 | 64 ± 3 | 15 ± 2 | 22 ± 2 | 17 ± 4 | 5.35 ± 1.27 | 5.54 ± 0.08 | 5.97 ± 0.42 | 0.41 ± 0.03 |
| | 5 – 10 | 55 ± 3 | 20 ± 1 | 24 ± 3 | 20 ± 5 | 0.28 ± 0.13 | 5.54 ± 0.06 | 4.67 ± 0.19 | 0.32 ± 0.01 |
| | 10 – 15 | 56 ± 2 | 20 ± 2 | 24 ± 2 | 11 ± 3 | 0.22 ± 0.12 | 5.49 ± 0.05 | 3.86 ± 0.11 | 0.26 ± 0.01 |
| | 15 – 20 | 53 ± 2 | 21 ± 3 | 26 ± 2 | 13 ± 3 | 0.09 ± 0.03 | 5.52 ± 0.04 | 3.1 ± 0.19 | 0.21 ± 0.01 |
| | 20 – 25 | 54 ± 3 | 20 ± 3 | 25 ± 2 | 17 ± 4 | 1.21 ± 1.12 | 5.52 ± 0.06 | 2.61 ± 0.17 | 0.18 ± 0.01 |
| | 25 – 30 | 55 ± 3 | 20 ± 3 | 24 ± 1 | 22 ± 3 | 0.05 ± 0.04 | 5.58 ± 0.08 | 2.34 ± 0.22 | 0.16 ± 0.01 |
| Bog | 0 – 5 | 60 ± 6 | 23 ± 6 | 17 ± 0 | 7 ± 3 | 12.41 ± 2.03 | 5.38 ± 0.10 | 12.78 ± 0.96 | 0.69 ± 0.07 |
| | 5 – 10 | 75 ± 2 | 19 ± 2 | 6 ± 1 | 121 ± 67 | 3.79 ± 1.16 | 5.51 ± 0.04 | 9.59 ± 4.26 | 0.54 ± 0.18 |
| | 10 – 15 | 80 ± 2 | 14 ± 3 | 6 ± 1 | 72 ± 44 | 1.01 ± 0.19 | 5.58 ± 0.02 | 11.47 ± 3.27 | 0.72 ± 0.14 |
| | 15 – 20 | 79 ± 3 | 15 ± 2 | 6 ± 1 | 119 ± 57 | 1.30 ± 0.26 | 5.53 ± 0.00 | 8.29 ± 2.67 | 0.5 ± 0.15 |
| | 20 – 25 | 82 ± 1 | 11 ± 2 | 7 ± 0 | 203 ± 63 | 2.52 ± 1.07 | 5.63 ± 0.02 | 5.82 ± 0.76 | 0.34 ± 0.07 |
| | 25 – 30 | 82 ± 5 | 11 ± 3 | 7 ± 2 | 129 ± 46 | 2.05 ± 0.58 | 5.69 ± 0.02 | 3.72 ± 0.72 | 0.2 ± 0.04 |

**Table S4**. Barcode identification for each of the samples analyzed. Raw data were deposited under the study accession number PRJNA384296 in the NCBI Sequence Read Archive (SRA). For 16S rRNA genes, primers used: F515 (5'-GTGCCAGCMGCCGCGGTAA-3’), R806 (5'-GGACTACVSGGGTATCTAAT-3’). For *pmoA* genes, primer set first round PCR (A189f/A682r) and second round multiplex PCR (A189f/A650r/mb661r): A189f (5’-GGNGACTGGGACTTCTGG-3’), A682r (5’-GAASGCNGAGAAGAASGC-3’), A650r (5’-ACGTCCTTACCGAAGGT-3’), mb661r (5’-CCGGMGCAACGTCYTTACC-3’).

| **Sample** | **Target gene** | **Soil – file name** | **Barcode** |
|---|---|---|---|
| B1-0 | 16S rRNA | fresh soil-bog- depth0cm-replicate-1 | GTCACA |
| B2-0 | 16S rRNA | fresh soil-bog- depth0cm-replicate-2 | TAGCAT |
| B3-0 | 16S rRNA | fresh soil-bog- depth0cm-replicate-3 | ACGTAC |
| B4-0 | 16S rRNA | fresh soil-bog- depth0cm-replicate-4 | TCAGAG |
| B1-5 | 16S rRNA | fresh soil-bog- depth5cm-replicate-1 | AGCTGA |
| B2-5 | 16S rRNA | fresh soil-bog- depth5cm-replicate-2 | CACAGT |
| B3-5 | 16S rRNA | fresh soil-bog- depth5cm-replicate-3 | AGAGTC |
| B4-5 | 16S rRNA | fresh soil-bog- depth5cm-replicate-4 | CGTATA |
| B1-10 | 16S rRNA | fresh soil-bog- depth10cm-replicate-1 | AGTCAG |
| B2-10 | 16S rRNA | fresh soil-bog- depth10cm-replicate-2 | CAGTCA |
| B3-10 | 16S rRNA | fresh soil-bog- depth10cm-replicate-3 | AGCTGA |
| B4-10 | 16S rRNA | fresh soil-bog- depth10cm-replicate-4 | GACTAG |
| B1-15 | 16S rRNA | fresh soil-bog- depth15cm-replicate-1 | ATATCG |
| B2-15 | 16S rRNA | fresh soil-bog- depth15cm-replicate-2 | CATGAC |
| B3-15 | 16S rRNA | fresh soil-bog- depth15cm-replicate-3 | AGTCAG |
| B4-15 | 16S rRNA | fresh soil-bog- depth15cm-replicate-4 | GAGATC |
| B1-20 | 16S rRNA | fresh soil-bog- depth20cm-replicate-1 | ATCGAT |
| B2-20 | 16S rRNA | fresh soil-bog- depth20cm-replicate-2 | CGATAT |
| B3-20 | 16S rRNA | fresh soil-bog- depth20cm-replicate-3 | ATATCG |
| B4-20 | 16S rRNA | fresh soil-bog- depth20cm-replicate-4 | GATCGA |
| B1-25 | 16S rRNA | fresh soil-bog- depth25cm-replicate-1 | ATGCTA |
| B2-25 | 16S rRNA | fresh soil-bog- depth25cm-replicate-2 | CGCGCG |
| B3-25 | 16S rRNA | fresh soil-bog- depth25cm-replicate-3 | ATCGAT |
| B4-25 | 16S rRNA | fresh soil-bog- depth25cm-replicate-4 | GTACAC |
| F1-0 | 16S rRNA | fresh soil-forest- depth0cm-replicate-1 | TACGTA |
| F2-0 | 16S rRNA | fresh soil-forest- depth0cm-replicate-2 | TATACG |
| F3-0 | 16S rRNA | fresh soil-forest- depth0cm-replicate-3 | ACTGCA |
| F4-0 | 16S rRNA | fresh soil-forest- depth0cm-replicate-4 | TCTCTC |
| F1-5 | 16S rRNA | fresh soil-forest- depth5cm-replicate-1 | ACACGT |
| F2-5 | 16S rRNA | fresh soil-forest- depth5cm-replicate-2 | AGTCAG |
| F3-5 | 16S rRNA | fresh soil-forest- depth5cm-replicate-3 | CGTATA |
| F4-5 | 16S rRNA | fresh soil-forest- depth5cm-replicate-4 | CAGTCA |
| F1-10 | 16S rRNA | fresh soil-forest- depth10cm-replicate-1 | ACGTAC |
| F2-10 | 16S rRNA | fresh soil-forest- depth10cm-replicate-2 | ATATCG |
| F3-10 | 16S rRNA | fresh soil-forest- depth10cm-replicate-3 | GACTAG |
| F4-10 | 16S rRNA | fresh soil-forest- depth10cm-replicate-4 | CATGAC |
| F1-15 | 16S rRNA | fresh soil-forest- depth15cm-replicate-1 | ACTGCA |
| F2-15 | 16S rRNA | fresh soil-forest- depth15cm-replicate-2 | ATCGAT |
| F3-15 | 16S rRNA | fresh soil-forest- depth15cm-replicate-3 | GAGATC |

| | | | |
|---|---|---|---|
| F4-15 | 16S rRNA | fresh soil-forest- depth15cm-replicate-4 | CGATAT |
| F1-20 | 16S rRNA | fresh soil-forest- depth20cm-replicate-1 | AGAGTC |
| F2-20 | 16S rRNA | fresh soil-forest- depth20cm-replicate-2 | ATGCTA |
| F3-20 | 16S rRNA | fresh soil-forest- depth20cm-replicate-3 | GATCGA |
| F4-20 | 16S rRNA | fresh soil-forest- depth20cm-replicate-4 | CGCGCG |
| F1-25 | 16S rRNA | fresh soil-forest- depth25cm-replicate-1 | AGCTGA |
| F2-25 | 16S rRNA | fresh soil-forest- depth25cm-replicate-2 | CACAGT |
| F3-25 | 16S rRNA | fresh soil-forest- depth25cm-replicate-3 | GTACAC |
| G1-0 | 16S rRNA | fresh soil-grass- depth0cm-replicate-1 | GTGTGT |
| G2-0 | 16S rRNA | fresh soil-grass- depth0cm-replicate-2 | GTGTGT |
| G3-0 | 16S rRNA | fresh soil-grass- depth0cm-replicate-3 | ATGCTA |
| G4-0 | 16S rRNA | fresh soil-grass- depth0cm-replicate-4 | TCGAGA |
| G1-5 | 16S rRNA | fresh soil-grass- depth5cm-replicate-1 | GACTAG |
| G2-5 | 16S rRNA | fresh soil-grass- depth5cm-replicate-2 | TACGTA |
| G3-5 | 16S rRNA | fresh soil-grass- depth5cm-replicate-3 | CACAGT |
| G4-5 | 16S rRNA | fresh soil-grass- depth5cm-replicate-4 | TCTCTC |
| G1-10 | 16S rRNA | fresh soil-grass- depth10cm-replicate-1 | GAGATC |
| G2-10 | 16S rRNA | fresh soil-grass- depth10cm-replicate-2 | TAGCAT |
| G3-10 | 16S rRNA | fresh soil-grass- depth10cm-replicate-3 | CAGTCA |
| G4-10 | 16S rRNA | fresh soil-grass- depth10cm-replicate-4 | ACACGT |
| G1-15 | 16S rRNA | fresh soil-grass- depth15cm-replicate-1 | GATCGA |
| G2-15 | 16S rRNA | fresh soil-grass- depth15cm-replicate-2 | TATACG |
| G3-15 | 16S rRNA | fresh soil-grass- depth15cm-replicate-3 | CATGAC |
| G4-15 | 16S rRNA | fresh soil-grass- depth15cm-replicate-4 | ACGTAC |
| G1-20 | 16S rRNA | fresh soil-grass- depth20cm-replicate-1 | GTACAC |
| G2-20 | 16S rRNA | fresh soil-grass- depth20cm-replicate-2 | TCAGAG |
| G3-20 | 16S rRNA | fresh soil-grass- depth20cm-replicate-3 | CGATAT |
| G4-20 | 16S rRNA | fresh soil-grass- depth20cm-replicate-4 | ACTGCA |
| G1-25 | 16S rRNA | fresh soil-grass- depth25cm-replicate-1 | GTCACA |
| G2-25 | 16S rRNA | fresh soil-grass- depth25cm-replicate-2 | TCGAGA |
| G3-25 | 16S rRNA | fresh soil-grass- depth25cm-replicate-3 | CGCGCG |
| G4-25 | 16S rRNA | fresh soil-grass- depth25cm-replicate-4 | AGAGTC |
| pmoA-B1-0 | *pmoA* | fresh soil-bog- depth0cm-replicate-1 | CACAGT |
| pmoA-B2-0 | *pmoA* | fresh soil-bog- depth0cm-replicate-2 | GACTAG |
| pmoA-B3-0 | *pmoA* | fresh soil-bog- depth0cm-replicate-3 | TACGTA |
| pmoA-B1-5 | *pmoA* | fresh soil-bog- depth5cm-replicate-1 | CAGTCA |
| pmoA-B2-5 | *pmoA* | fresh soil-bog- depth5cm-replicate-2 | GAGATC |
| pmoA-B3-5 | *pmoA* | fresh soil-bog- depth5cm-replicate-3 | TAGCAT |
| pmoA-B1-10 | *pmoA* | fresh soil-bog- depth10cm-replicate-1 | CATGAC |
| pmoA-B2-10 | *pmoA* | fresh soil-bog- depth10cm-replicate-2 | GATCGA |
| pmoA-B3-10 | *pmoA* | fresh soil-bog- depth10cm-replicate-3 | TATACG |
| pmoA-B1-15 | *pmoA* | fresh soil-bog- depth15cm-replicate-1 | CGATAT |
| pmoA-B2-15 | *pmoA* | fresh soil-bog- depth15cm-replicate-2 | GTACAC |
| pmoA-B3-15 | *pmoA* | fresh soil-bog- depth15cm-replicate-3 | TCAGAG |
| pmoA-B1-20 | *pmoA* | fresh soil-bog- depth20cm-replicate-1 | CGCGCG |
| pmoA-B2-20 | *pmoA* | fresh soil-bog- depth20cm-replicate-2 | GTCACA |
| pmoA-B3-20 | *pmoA* | fresh soil-bog- depth20cm-replicate-3 | TCGAGA |
| pmoA-B1-25 | *pmoA* | fresh soil-bog- depth25cm-replicate-1 | CGTATA |
| pmoA-B2-25 | *pmoA* | fresh soil-bog- depth25cm-replicate-2 | GTGTGT |

| | | | |
|---|---|---|---|
| pmoA-B3-25 | *pmoA* | fresh soil-bog- depth25cm-replicate-3 | TCTCTC |
| pmoA-F1-0 | *pmoA* | fresh soil-forest- depth0cm-replicate-1 | ACACGT |
| pmoA-F2-0 | *pmoA* | fresh soil-forest- depth0cm-replicate-2 | ATATCG |
| pmoA-F3-0 | *pmoA* | fresh soil-forest- depth0cm-replicate-3 | CGATAT |
| pmoA-F1-5 | *pmoA* | fresh soil-forest- depth5cm-replicate-1 | ACGTAC |
| pmoA-F2-5 | *pmoA* | fresh soil-forest- depth5cm-replicate-2 | ATCGAT |
| pmoA-F3-5 | *pmoA* | fresh soil-forest- depth5cm-replicate-3 | CGCGCG |
| pmoA-F1-10 | *pmoA* | fresh soil-forest- depth10cm-replicate-1 | ACTGCA |
| pmoA-F2-10 | *pmoA* | fresh soil-forest- depth10cm-replicate-2 | ATGCTA |
| pmoA-F3-10 | *pmoA* | fresh soil-forest- depth10cm-replicate-3 | CGTATA |
| pmoA-F1-15 | *pmoA* | fresh soil-forest- depth15cm-replicate-1 | AGAGTC |
| pmoA-F2-15 | *pmoA* | fresh soil-forest- depth15cm-replicate-2 | CACAGT |
| pmoA-F3-15 | *pmoA* | fresh soil-forest- depth15cm-replicate-3 | GACTAG |
| pmoA-F1-20 | *pmoA* | fresh soil-forest- depth20cm-replicate-1 | AGCTGA |
| pmoA-F2-20 | *pmoA* | fresh soil-forest- depth20cm-replicate-2 | CAGTCA |
| pmoA-F3-20 | *pmoA* | fresh soil-forest- depth20cm-replicate-3 | GAGATC |
| pmoA-F1-25 | *pmoA* | fresh soil-forest- depth25cm-replicate-1 | AGTCAG |
| pmoA-F2-25 | *pmoA* | fresh soil-forest- depth25cm-replicate-2 | CATGAC |
| pmoA-F3-25 | *pmoA* | fresh soil-forest- depth25cm-replicate-3 | GATCGA |
| pmoA-G1-0 | *pmoA* | fresh soil-grass- depth0cm-replicate-1 | GTACAC |
| pmoA-G2-0 | *pmoA* | fresh soil-grass- depth0cm-replicate-2 | TCAGAG |
| pmoA-G3-0 | *pmoA* | fresh soil-grass- depth0cm-replicate-3 | AGAGTC |
| pmoA-G1-5 | *pmoA* | fresh soil-grass- depth5cm-replicate-1 | GTCACA |
| pmoA-G2-5 | *pmoA* | fresh soil-grass- depth5cm-replicate-2 | TCGAGA |
| pmoA-G3-5 | *pmoA* | fresh soil-grass- depth5cm-replicate-3 | AGCTGA |
| pmoA-G1-10 | *pmoA* | fresh soil-grass- depth10cm-replicate-1 | GTGTGT |
| pmoA-G2-10 | *pmoA* | fresh soil-grass- depth10cm-replicate-2 | TCTCTC |
| pmoA-G3-10 | *pmoA* | fresh soil-grass- depth10cm-replicate-3 | AGTCAG |
| pmoA-G1-15 | *pmoA* | fresh soil-grass- depth15cm-replicate-1 | TACGTA |
| pmoA-G2-15 | *pmoA* | fresh soil-grass- depth15cm-replicate-2 | ACACGT |
| pmoA-G3-15 | *pmoA* | fresh soil-grass- depth15cm-replicate-3 | ATATCG |
| pmoA-G1-20 | *pmoA* | fresh soil-grass- depth20cm-replicate-1 | TAGCAT |
| pmoA-G2-20 | *pmoA* | fresh soil-grass- depth20cm-replicate-2 | ACGTAC |
| pmoA-G3-20 | *pmoA* | fresh soil-grass- depth20cm-replicate-3 | ATCGAT |
| pmoA-G1-25 | *pmoA* | fresh soil-grass- depth25cm-replicate-1 | TATACG |
| pmoA-G2-25 | *pmoA* | fresh soil-grass- depth25cm-replicate-2 | ACTGCA |
| pmoA-G3-25 | *pmoA* | fresh soil-grass- depth25cm-replicate-3 | ATGCTA |

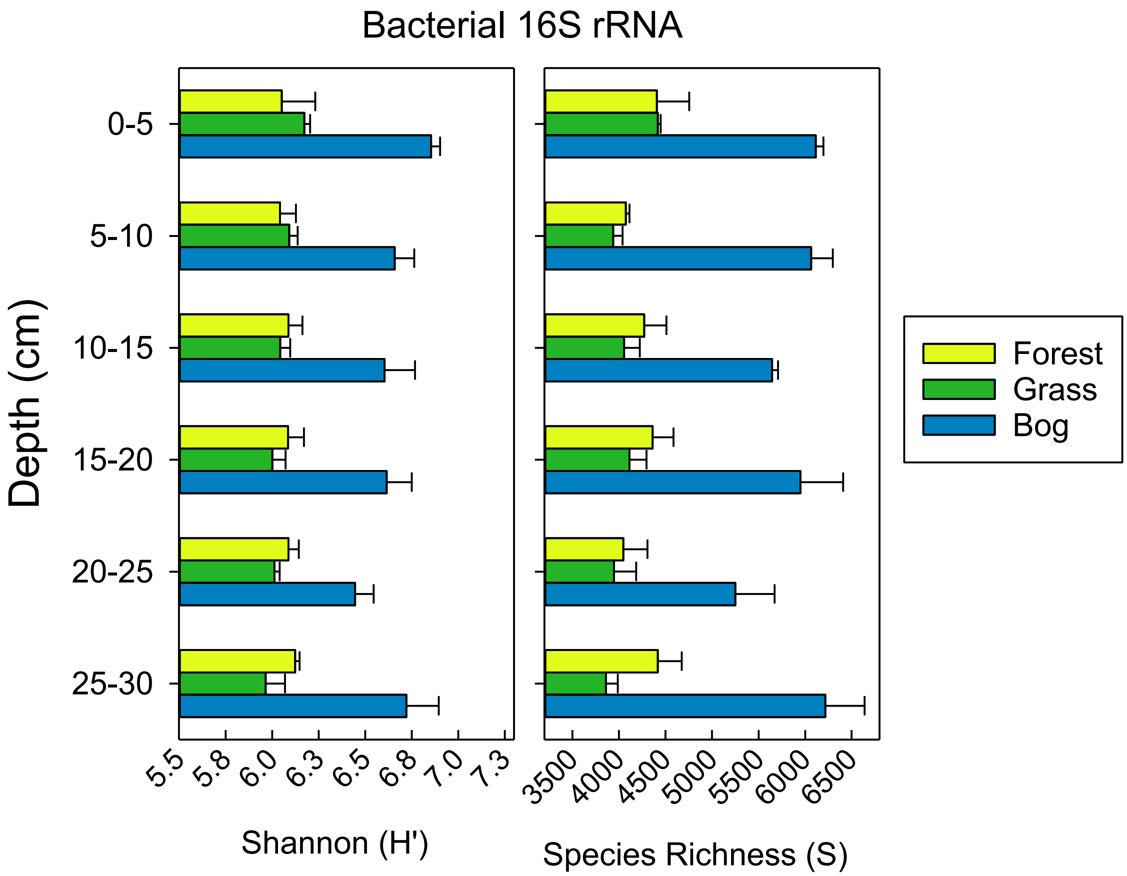

**Figure S1**. Bacterial 16S rRNA gene diversity measured as Shannon Diversity (H') and Richness (S) (n=4, means ± standard errors).

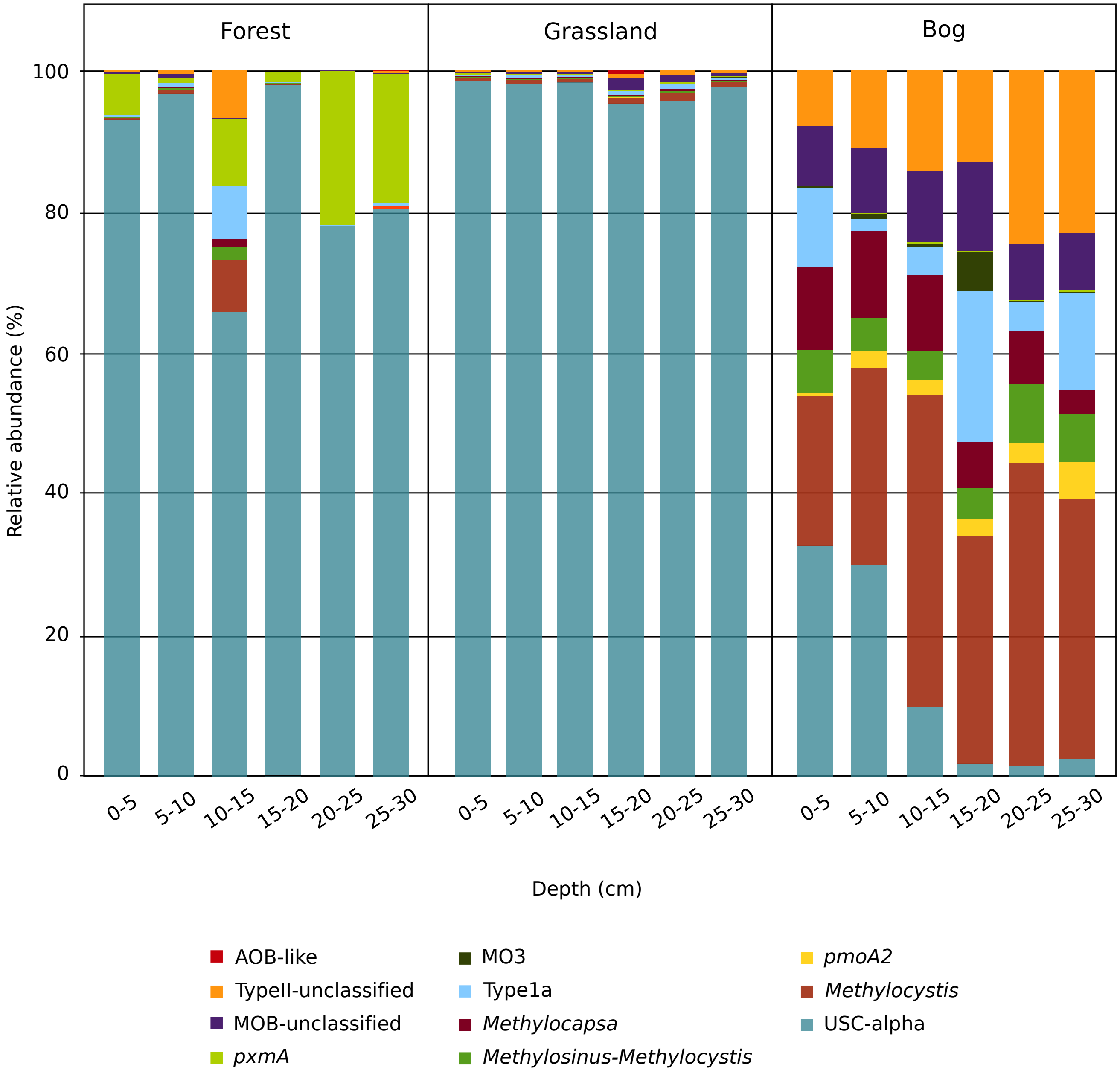

**Figure S2.** Relative abundance of *pmoA* genes of the dominant methanotroph groups.

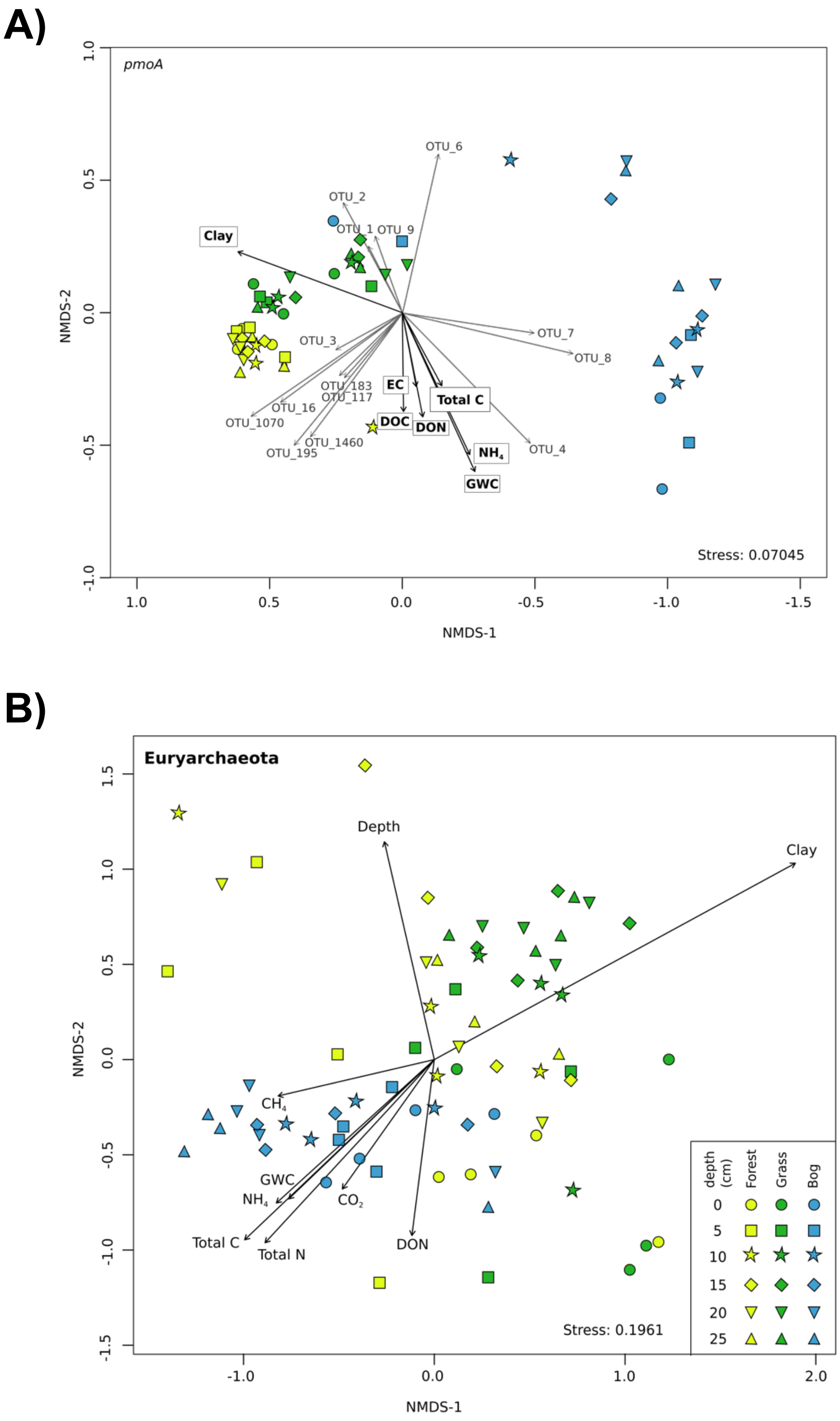

**Figure S3**. NMDS ordination of *pmoA* (A) and euryarchaeota (B) communities based on the Bray–Curtis dissimilarity of community composition. Shape indicates depth and sites are colored according to the soil typel. The arrows indicate the direction at which the environmental vectors fit the best (using the *envfit* function) onto the NMDS ordination space. Abbreviations: DOC, dissolved organic carbon; DON, dissolved organic nitrogen; EC, electrical conductivity; GWC, gravimetric water content; $NH_4$, ammonium.

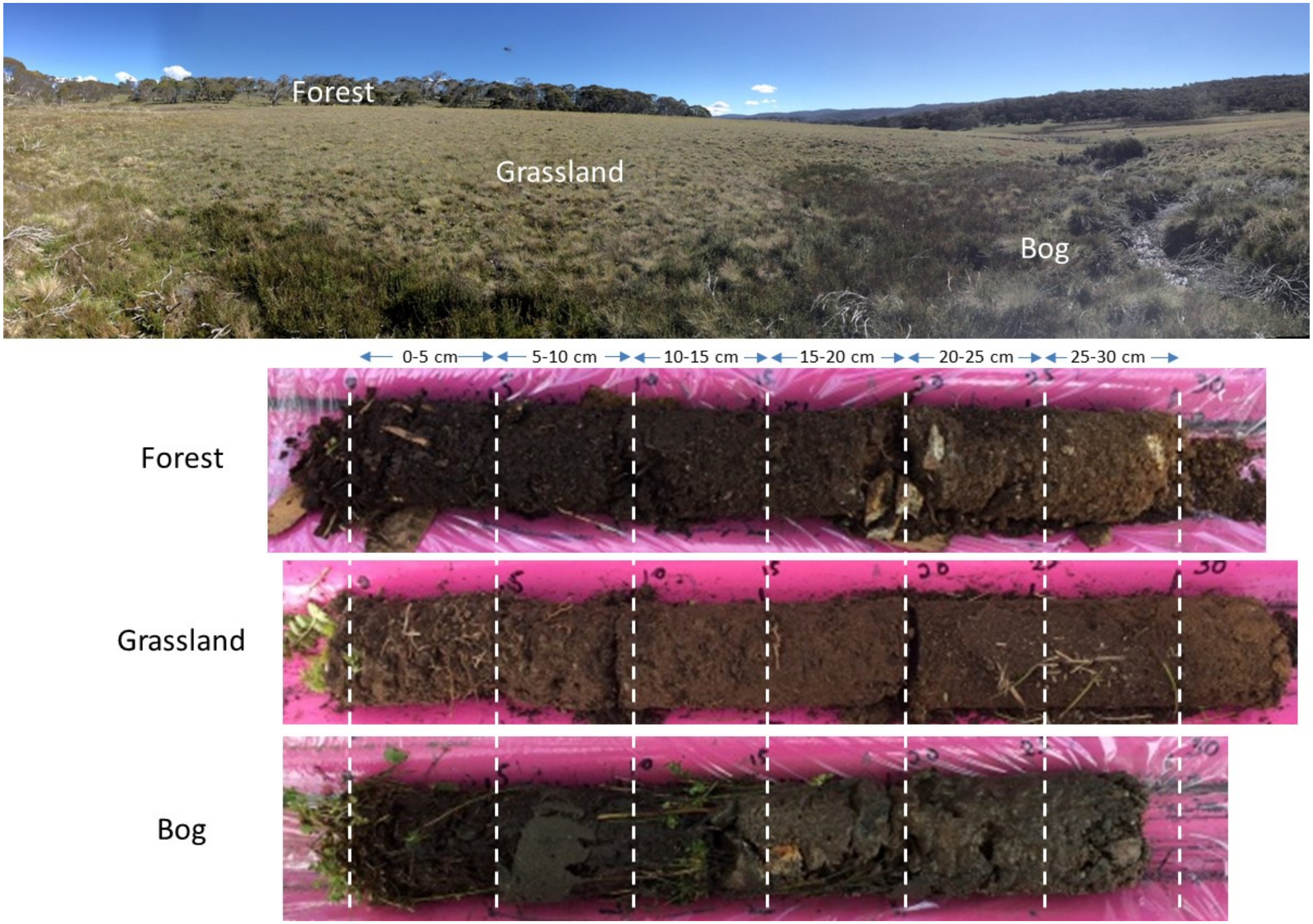

**Figure S4**. Soil/Vegetation gradient (Forest-Grassland-Bog) located in Snowy Mountains region near Kosciuszko National Park, NSW, Australia. Soil/vegetation types shown above, below are three soil cores from each of the soil/vegetation types.
